# Super-resolution vibrational imaging using expansion stimulated Raman scattering microscopy

**DOI:** 10.1101/2021.12.22.473713

**Authors:** Lixue Shi, Aleksandra Klimas, Brendan Gallagher, Zhangyu Cheng, Feifei Fu, Piyumi Wijesekara, Yupeng Miao, Xi Ren, Yongxin Zhao, Wei Min

## Abstract

Stimulated Raman scattering (SRS) microscopy is an emerging technology that provides high chemical specificity for endogenous biomolecules and can circumvent common constraints of fluorescence microscopy including limited capabilities to probe small biomolecules and difficulty resolving many colors simultaneously due to spectral overlap. However, the resolution of SRS microscopy remains governed by the diffraction limit. To overcome this, we describe a new technique called Molecule Anchorable Gel-enabled Nanoscale Imaging of Fluorescence and stImulatEd Raman Scattering microscopy (MAGNIFIERS), that integrates SRS microscopy with expansion microscopy (ExM). ExM is a powerful strategy providing significant improvement in imaging resolution by physical magnification of hydrogel-embedded preserved biological specimens. MAGNIFIERS offers chemical-specific nanoscale imaging with sub-50 nm resolution and has scalable multiplexity when combined with multiplex Raman probes and fluorescent labels. We used MAGNIFIERS to visualize nanoscale features in a label-free manner with C-H vibration of proteins, lipids and DNA in a broad range of biological specimens, from mouse brain, liver and kidney to human lung organoid. In addition, we applied MAGNIFIERS to track nanoscale features of protein synthesis in protein aggregates using metabolic labeling of small metabolites. Finally, we used MAGNIFIERS to demonstrate 8-color nanoscale imaging in an expanded mouse brain section. Overall, MAGNIFIERS is a valuable platform for super-resolution label-free chemical imaging, high-resolution metabolic imaging, and highly multiplexed nanoscale imaging, thus bringing SRS to nanoscopy.

## Introduction

While fluorescence microscopy has been the predominant imaging modality in biophotonics, it faces fundamental limitations for many advanced imaging applications. Going beyond fluorescence, stimulated Raman scattering (SRS) microscopy is emerging as a powerful modality for illuminating otherwise invisible chemical bonds of biomolecules in complex biological contexts^1, 2^. In contrast to fluorescence imaging, SRS generates high chemical specificity via probing vibrational transitions, enabling label-free detection of vital biomolecules of proteins, lipids, nucleic acids, etc^1, 3^. Moreover, integration of SRS with vibrational tags renders a paradigm of bioorthogonal chemical imaging for studying metabolic activities in cells and tissues^2, 4, 5^. The tiny size and good biocompatibility of vibrational tags, such as stable isotopes and triple bonds, enable non-perturbative tracing of small molecules^4^, which is difficult to achieve with fluorescence because of the bulky fluorophores. Furthermore, by virtue of much narrower vibrational peaks (~10 cm^-1^) compared to fluorescence peaks (~500 cm^-1^), Raman imaging offers scalable multiplexity and breaks the ‘palette number barrier’ of fluorescence, which can typically only image 4-5 colors simultaneously due to the overlap of inherently broad fluorescence spectra. In this regard, development of rainbow-like Raman dyes leverages the sensitivity of SRS and opens the door for supermultiplexed vibrational imaging^6–8^. In particular, by combining electronic pre-resonance spectroscopy with stimulated Raman scattering (SRS) microscopy (i.e., epr-SRS), the Raman cross sections of electronically coupled vibrational modes in light-absorbing dyes can be enhanced by 10^13^-fold^6, 9^. Owing to this drastic enhancement, we achieved nanomolar sensitivity of Raman-active dyes (such as commercial far-red fluorescent dyes and specially-designed MARS probes), and demonstrated epr-SRS imaging of specific proteins^6, 10, 11^.

Collectively, SRS imaging circumvents the fundamental bottlenecks of fluorescence microscopy and provides unique benefits including label-free chemical imaging of endogenous biomolecules^1, 12^, nonperturbative mapping of small metabolites^4^, and simultaneous profiling of a large group of specific biomarkers^6, 11^. Yet, being an optical technique that uses near infrared or visible light, SRS microscopy is still constrained to diffraction-limited spatial resolution (~300 nm). Thus, many crucial nanoscale organizations in cells and tissues cannot be resolved by the current iteration of SRS microscopy. Despite extensive efforts in extending Raman imaging beyond the diffraction limit, sophisticated instrumentation is required and only moderate resolution improvement (typically less than 2-fold) has been achieved, resulting in limited utilization in biological specimens^13–18^. Hence, vibrational-based nanoscale imaging approaches are still underexplored territory in biomedical research.

Over the last half decade, a sample-oriented nanoscale imaging technique known as expansion microscopy (ExM) has emerged as a non-optical means of achieving effective resolutions beyond the diffraction limit. ExM is a powerful technique for nanoscale optical imaging through physical magnification of tissue samples embedded in a water-swellable polymer gel^19^. ExM was initially developed for fluorescence microscopy with a ~4.5× expansion in each dimension in pure water and quickly evolved to different variants with widespread use in biology and pathology^20–24^. Conceptually, the principle of ExM could be applied to overcome the diffraction limit of SRS imaging. However, common ExM variants^22, 23, 25^ rely on strong protease digestion to facilitate expansion, thereby destroying endogenous proteins. On the other hand, ExM variants that retain proteins after expansion^26–28^ are more compatible with SRS imaging, but they require special fixation protocols and are only validated on culture cells and mouse brain tissues. A very recent technique called VISTA was developed for label-free chemical imaging of proteins on expanded mouse brain tissues using a protein-retention ExM variant^29^, but it inherits the critical limitation of this ExM variant on tissue compatibility and resolution gain.

Recently, a new ExM framework has emerged that offers larger expansion factors along with broad biomolecule retention and tissue compatibility^30^, capable of expanding a multitude of tissue types with standard fixation, from PFA-fixed mouse brain and human lung organoid to an array of clinically relevant formalin-fixed paraffin-embedded (FFPE) human tissues which have proved challenging to expand in the past. By combining SRS imaging with this framework, we lay out an approach named <u>M</u>olecule <u>A</u>nchorable <u>G</u>el-enabled <u>N</u>anoscale <u>I</u>maging of <u>F</u>luorescence and st<u>I</u>mulat<u>E</u>d <u>R</u>aman <u>S</u>cattering microscopy (MAGNIFIERS). Harnessing the superior retention of biomolecular contents and general sample compatibility across multiple tissue types and fixation methods of the new ExM framework, the MAGNIFIERS approach provides excellent reversible linear expansion up to ~7.2-fold providing lateral resolutions of ~41 nm (298 nm/7.2=41.4 nm) and axial resolutions of ~194 nm (1400 nm/7.2=194 nm), with great flexibility to couple vibrational metabolic labeling and Raman dye staining.

MAGNIFIERS revitalizes a full spectrum of utilities of SRS imaging into the sub-diffraction regime (Fig. 1a). Specifically, we show that MAGNIFIERS supports label-free nanoscale imaging of endogenous biomolecules including proteins, lipids, and DNA. We further demonstrate its power for high-resolution metabolic imaging of protein aggregates in cells labeled with deuterium-substituted amino acids. Finally, we use MAGNIFIERS to map the nanoscale locations of 8 different markers in mouse brain tissue in one shot by applying multiplex Raman labels and conventional fluorescent probes.

**Fig. 1.**
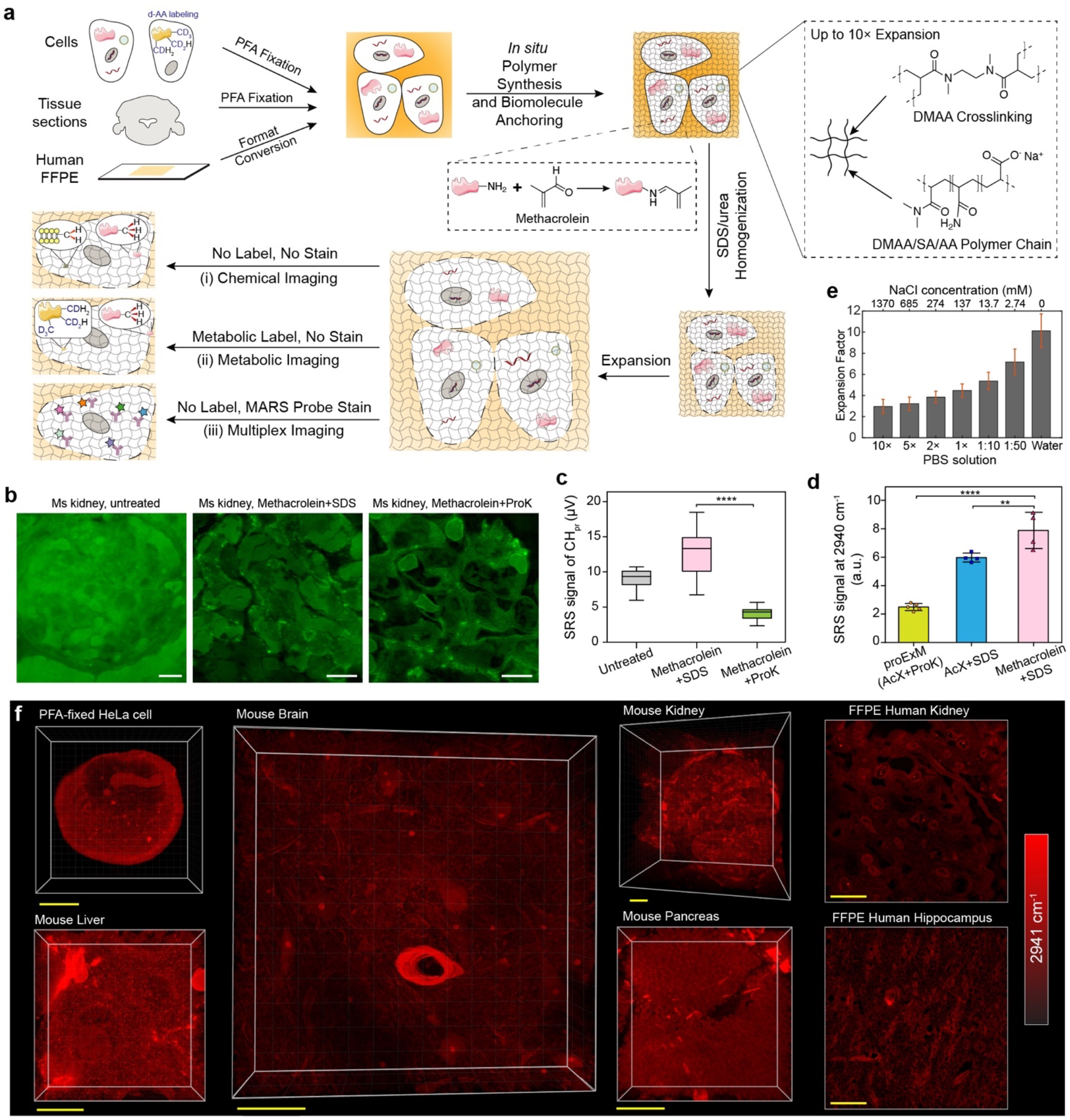
Integration of SRS with an optimized ExM protocol (MAGNIFY). **a**, Schematics of the workflow of MAGNIFIERS and gel chemistry of MAGNIFY. During *in situ* polymer synthesis, methacrolein was added as part of the monomer solution to efficiently attach biomolecules to the growing matrix. The gel was formed from polymerization of cocktails of the monomers acrylamide (AA, nonionic monomer), sodium acrylate (SA, ionic monomer) and *N,N*-dimethylacrylamide (DMAA, nonionic self-cross-linker. Afterwards, the gel-embedded specimens were treated with sodium dodecyl sulfate (SDS)/urea solution in 80-90 °C for at least one hour for homogenization. PFA, paraformaldehyde. **b**, SRS images of unmixed CH_protein_ signal of the glomerulus in the mouse kidney tissues. (left) untreated, (middle) expanded with methacrolein-linking and SDS/urea homogenization, (right) expanded with methacrolein-linking and Proteinase K digestion. **c**, Quantification of protein contents for untreated and expanded samples as shown in (**b**). SRS signals were plotted as mean±s.d. (n=29, 25, 25 regions of interest (ROIs)). In the box plot, the center indicates the median; the bottom and top edges of the box indicate the 25^th^ and 75^th^ percentiles, respectively; the whiskers extend to the minimum and maximum data points. Absolute signal in the untreated sample is converted with a volume dilution factor of 4.3^3^= ~80-fold after expansion. Two-tailed unpaired *t*-test, \*\*\*\**P*<0.0001, *t*=13. **d**, Comparison of protein retention on the proExM protocol with the MAGNIFIERS protocol. SRS signals were plotted as mean±s.d. (n=5, 4, 4 ROIs). One-way ANOVA followed by Bonferroni’s *post hoc* test using the ‘Methacrolein+SDS’ as the control column, \*\*\*\**P*<0.0001 and \*\**P*=0.0085. **e**, Measurement of expansion factors (mean±s.d.) in PBS buffers with different salt concentrations. Corresponding salt concentrations of used PBS buffer solution are labeled on the up X axis. Values of n are provided in Supplementary Table 2. **f**, 3D-rendered SRS images of CH_3_ peak at 2941 cm^-1^ of PFA-fixed HeLa cell, mouse brain, liver, kidney and pancreas tissues, and FFPE human kidney and brain hippocampus tissue. Yellow scale bars are not corrected for the expansion factor. Scale bars, 10 μm in (**b**); 50 μm (postexpansion) in (**f**).

## Results

### Integration of SRS with an optimized biomolecule-retention ExM protocol (MAGNIFY)

To fully demonstrate the potential of expansion SRS imaging, we chose a new ExM strategy that achieves a favorable balance between biomolecule retention, expansion factor, and sample generality^30^. The new method is called Molecule Anchorable Gel-enabled Nanoscale In-situ Fluorescence MicroscopY (MAGNIFY)^30^. To demonstrate its utility for SRS imaging, we first quantified the protein retention rate by imaging CH_3_ stretching vibration of proteins at 2941 cm^-1^ in paraformaldehyde (PFA)-fixed mouse kidney tissue pre- and post-expansion (Fig. 1b). Remarkably, we found a near 100% protein retention rate using the MAGNIFY approach (Fig. 1b-c), substantially higher than that of protein-retention expansion microscopy using 6-((acryloyl)amine) hexanoic acid (AcX) as an anchoring agent (i.e. proExM^22^, Fig. 1d and Supplementary Fig.1), ensuring optimal sensitivity and effective resolution. The reasons for the optimized biomolecule retention rate are twofold. First, methacrolein was employed as part of the monomer solution to efficiently attach biomolecules to the growing matrix during the *in situ* polymerization step, eliminating the need of stepwise reactions of anchoring and polymerization (Fig. 1a). Second, rather than strong protease digestion, the gel-embedded specimens were treated with a denaturant-rich solution in 80-90 °C for homogenization to ensure maximum retention of biomolecules in the gel.

MAGNIFIERS relied on a reformulated hydrogel (Fig. 1a) which allows isotropic physical expansion up to 10-fold in a single round^30^. To theoretically estimate the improved spatial precision, we calibrated the resolution of the current SRS microscopy by imaging single 100-nm polystyrene beads at 3056 cm^-1^ (Supplementary Fig. 2). Cross-section profiles fitted by the Gaussian function indicated the full width at half maximum (FWHM) of lateral direction of 298 nm (Supplementary Fig. 2a). Thus, effective resolution of ~30 nm is possible given an expansion factor (EF) of around 10 (Fig. 1e). Unless stated otherwise, the reformulated hydrogel was used for expanding all biological specimens.

MAGNIFIERS is broadly applicable to a wide range of specimens (Supplementary Fig. 3), as it is compatible with common fixation methods and multiple tissue types, including mechanically tough tissues such as kidney and liver. For instance, SRS imaging of CH_3_ stretching at 2941 cm^-1^ visualized rich three-dimensional (3D) structural information in cultured cells and mouse tissues such as the brain, liver, kidney, and pancreas (Fig. 1f, Supplementary Video 1), thanks to the intrinsic 3D sectioning capability from the two-photon excitation nature of SRS. In addition, MAGNIFIERS is suitable for human organoids (Supplementary Fig. 3), a favorable model system for human biology and medicine. Going beyond freshly preserved cells and tissues, archival samples such as FFPE sections of human brain, kidney and spleen, which is a popular form of preparation for biopsy specimens that aids in clinical examination and pathology research, are also compatible with MAGNIFIERS (Fig. 1f, Supplementary Fig. 3).

### Label-free visualization of biological and pathological ultrastructures

Encouraged by the near 100% protein retention, we envision it is advantageous for unveiling nanoscale features without the need to label or stain with fluorescent dyes. In standard SRS microscopy, C-H stretching region at ~2800-3050 cm^-1^ is desirable to image endogenous species^3^. However, the expanded sample might exhibit O-H background (~2900-3700 cm^-1^) from the water (typically >90% by volume) in the sample and C-H background from the gel matrix itself, thus potentially overwhelming the signal of interest. To study this, we performed hyperspectral SRS analysis of expanded FFPE human kidney tissues (Fig. 2a, Supplementary Fig. 4a). We chose FFPE tissue as a test because of its simple Raman spectrum, as protein is the major component and most lipids are removed in deparaffinization before expansion (Supplementary Fig. 4b). By analyzing the spectra of a tissue region and a blank gel region (Fig. 2a, left), we observed strong C-H signals on the tissue area but only a background from the tail of water’s O-H vibration in the blank gel region (Fig. 2a, right). The absence of C-H peaks in the blank gel suggests that the C-H signal from the gel matrix is actually negligible, likely due to the low concentration of the hydrogel. After subtracting the water background, which is broad and smooth, the spectral shape recapitulates a typical CH_3_ mode of proteins with a major peak at ~2941 cm^-1^ (Fig. 2b), consistent with the fact that proteins are the major anchored biomolecules in FFPE tissue. Based on our spectral analyses, we confirmed that the polymer gel is “Raman transparent” in the C-H region and that the leak of the O-H background can be readily subtracted, validating using the CH_3_ vibration at 2941 cm^-1^ for imaging total proteins in expanded specimens.

**Fig. 2.**
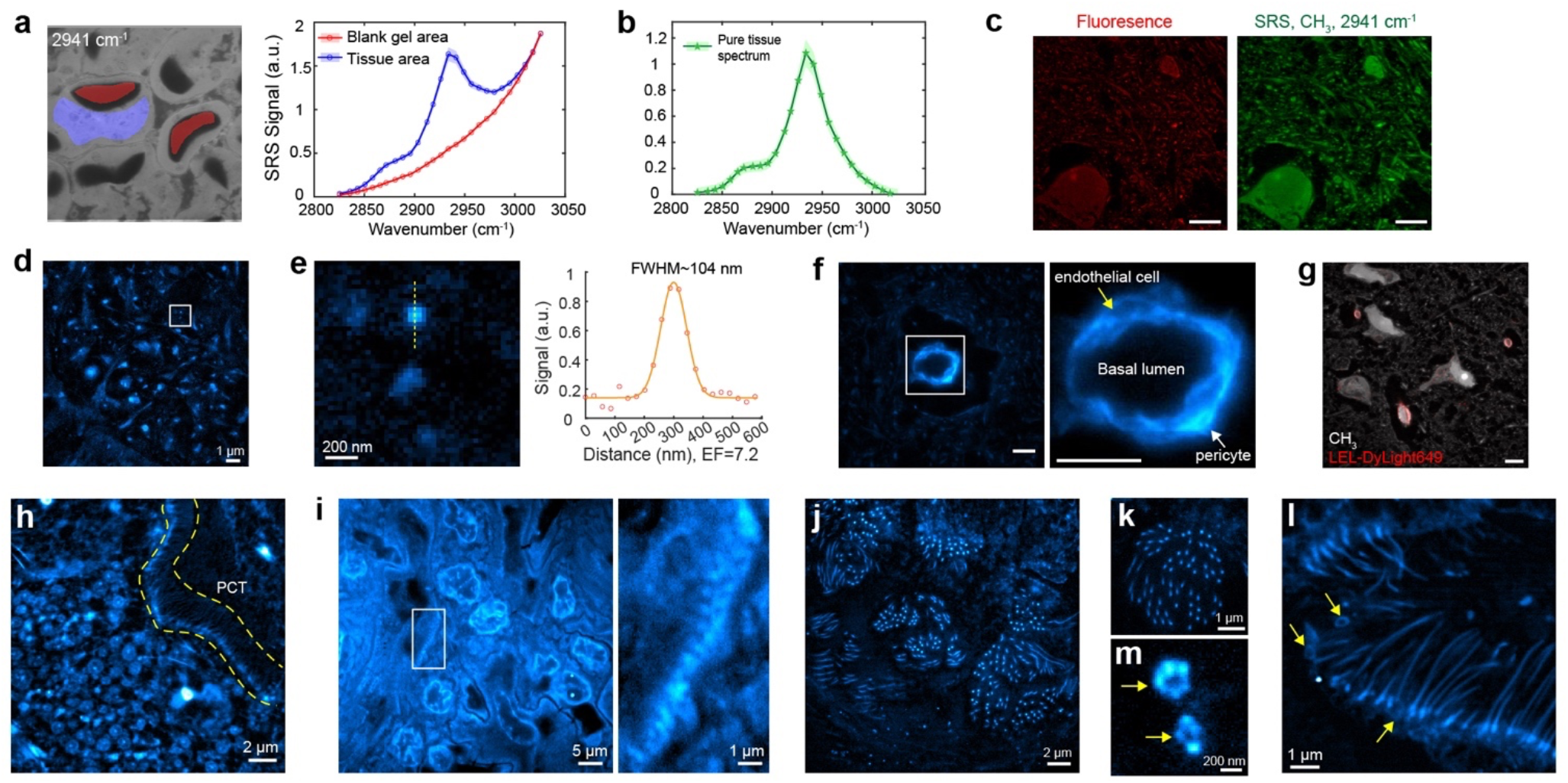
MAGNIFIERS visualizes ultrafine structures of biological samples via label-free protein imaging. **a**, Hyperspectral SRS imaging of the C-H stretching region on expanded FFPE human kidney. Left, masks of tissue area (red) and blank gel area (blue) for spectral analysis labeled on the SRS image of 2941 cm^-1^. Right, SRS spectra of two masked areas. Single-frequency SRS images are in Supplementary Fig. 4a. **b**, Background-removed SRS spectrum of expanded FFPE human kidney. **c**, SRS CH_3_ image at 2941 cm^-1^ and fluorescence image of Alexa Fluor 555 NHS ester labeled mouse brain tissue. **d**, CH_3_ image of extended mouse brain in 1/50× PBS (7.2-fold expansion) with a 1.2 NA objective. **e**, Zoom into the region outlined by the white box in (**d**). Right, line profile of a subdiffraction-limited spot outlined by the yellow line. **f**, Fine structures of blood-brain barrier in mouse brain tissue. Single endothelial cells and a pericyte were visualized. Images were collected using a 1.05 NA objective in 1× PBS (4.5-fold expansion). Right, zoom into the region outlined by the white box. **g**, Blood vessels and capillaries stand out in the protein channel, confirmed by lectin staining. Expanded mouse brain tissue labeled with *Lycopersicon Esculentum* lectin (LEL)-DyLight 649 and imaged with a 1.05 NA objective in 1× PBS (4.5-fold expansion). **h**, SRS protein image in expanded mouse kidney. PCT, proximal convoluted tubule. Yellow region highlights the cilia on the surface of PCT. **i**, SRS protein image reveals periodic podocyte foot processes in expanded FFPE human kidney tissue. Right, zoom into the region outlined by the white box. Images were collected using a 1.2 NA objective in 1 × PBS (2.3-fold expansion). **j**-**m**, Flower-like cilia structure (**j, k**) and the ring structure at the base of cilia (**l, m**) in the expanded human lung organoid were visualized by the protein channel at 2941 cm^-1^. Yellow arrows in (**k, m**) highlight the basal bodies. Images were collected with a 1.05 NA objective, (**j, l**) in 1× PBS (4.5-fold expansion) and (**k, m**) in 1/50× PBS (7.2-fold expansion). Scale bars, 200 nm in (**e, m**); 1 μm in (**d, i** (right), **k, l**); 2 μm in (**f, h, j**); 5 μm in (**c, g, i** (left)).

It is valuable to compare the protein distribution pattern from SRS imaging of CH_3_ vibration at 2941 cm^-1^ to that from fluorescence imaging of stained total protein content^31, 32^. By performing SRS and fluorescence imaging on an expanded mouse brain section stained with N-hydroxysuccidimidyl (NHS) ester-dye conjugates, we observed similar patterns (Fig. 2c). This morphological agreement between SRS and fluorescence images corroborates our prior spectral analysis.

The effective resolution is jointly determined by the objective’s numerical aperture (NA) and the achievable EF, which can be controlled by osmotic pressure of the imaging solution (Fig. 1e). Given a practical EF of 7.2 with good signal (Fig. 2d), an effective resolution of ~298/7.2=41 nm is feasible. With this improved resolution, we resolved sub-diffraction-limited dots on expanded mouse brain tissue, proved by a fitted FWHM of 104 nm (Fig. 2e).

MAGNIFIERS can unveil a broad range of biological or pathological ultrastructure features solely via label-free protein imaging. First, the microscopic view of blood-brain barrier in mouse brain tissue, including single endothelial cells and a pericyte, was visualized (Fig. 2f), supported by the co-staining with lectins as a blood vessel marker (Fig. 2g, Supplementary Fig. 4c). Benefiting from good sample compatibility of MAGNIFY, we further revealed interesting nanoscale architectures in both PFA-fixed mouse kidney and FFPE human kidney specimens. For instance, cilia were visualized at the inner layer of the proximal convoluted tubule (PCT) in mouse kidney (Fig. 2h). In addition, we observed sub-diffraction-limited structures of interdigitated foot processes of podocytes in the glomeruli of human (Fig. 2i) and mouse kidney tissues (Supplementary Fig. 5a-e), which is a clinical metric for many nephrotic kidney diseases such as kidney minimal change disease^33^. Moreover, in expanded human bronchial epithelial cells-derived lung organoids, flower-like cilia architectures were faithfully elucidated, projecting from the apical epithelial surface (Fig. 2j-k, Supplementary Fig. 5f, Supplementary Video 2). Intracellular structures and patterns of surrounding apoptotic bodies were also clearly observed (Supplementary Fig. 5g-h). Remarkably, at the basal foot of cilia, the ring structure of ciliary base bodies, which are composed of a 9+0 microtubule arrangement and have an inner diameter of around 200 nm^34^, were precisely identified (Fig. 2l-m, yellow arrows).

### Label-free nanoscale imaging of chemical compositions

The MAGNIFY framework is known to retain biomolecules other than proteins such as lipids and nucleic acids^30^. Thus, MAGNIFIERS holds great promise for label-free chemically-defined nanoscale imaging in a comprehensive view. Yet, the uncharacterized issue is whether SRS can detect these molecules with sufficient contrast. Prior SRS spectral studies have dissected the C-H signals into contributions from proteins (~2931 cm^-1^), lipids (~2854 cm^-1^) and DNA (~2956 cm^-1^) through linear spectral unmixing^12^. We started by applying this strategy to analyze expanded human lung organoids with an EF of 4.5. In the unmixed images which are labeled as CH_Pr_, CH_L_ and CH_DNA_ (Fig. 3a-b), we observed the presence of lipids in small-size extracellular vesicles^35^ (EVs, secreted membrane vesicles), some of which also contain CH_DNA_ signals. We next performed a hyperspectral analysis of the entire C-H region (Supplementary Fig. 6a). By examining the pixel-averaged SRS spectrum of all small EVs (i.e. exosomes) in the region of interest, a distinct side peak around 2865 cm^-1^ appeared (Fig. 3c, Supplementary Fig. 6b), in agreement with the peak signature of CH_2_ vibration from lipids. Exosomes contain high levels of cholesterol, sphingomyelin and ceramide, similar to the composition of detergent-resistant lipid rafts^36^. As a control, the SRS spectrum of the cell predominantly reflects the chemical footprint of proteins (Fig. 3c, blue). Furthermore, we generated the Raman spectrum of a single DNA-containing EV (pink arrowed in Fig. 3b) which displayed another pronounced shoulder peak of ~2955 cm^-1^ originating from DNA as we anticipated (Fig. 3d, Supplementary Fig. 6c). This observation is in line with prior reports demonstrating that EVs contain proteins, lipids and nucleic acids^35, 37^.

**Fig. 3.**
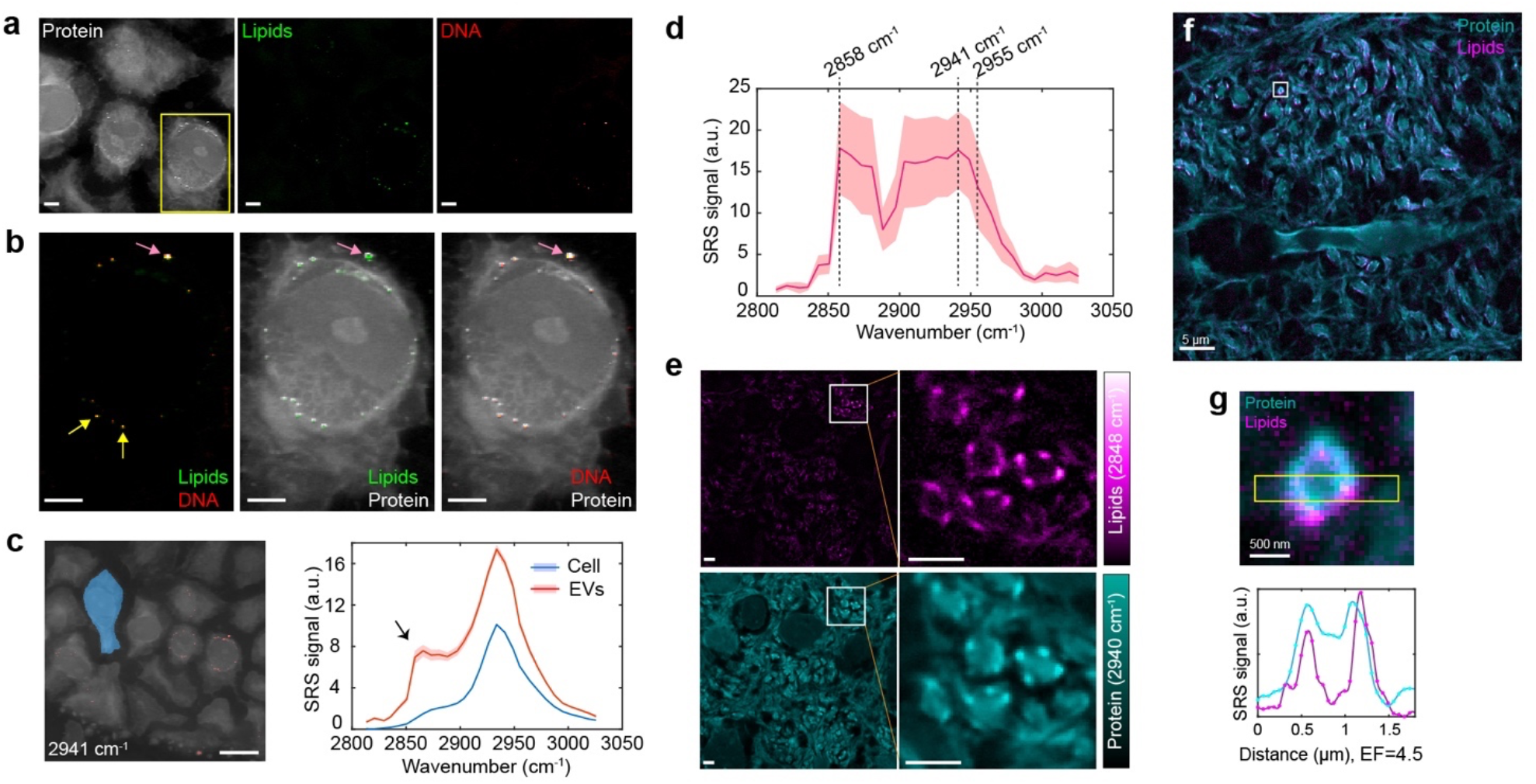
Label-free nanoscale imaging of chemical compositions. **a**, Spectrally unmixed C-H channels for protein, lipids and DNA in expanded human lung organoid. **b**, Two-color overlayed images of zoomin areas outlined by the yellow box in (**a**). Arrows indicate EVs containing lipids and DNA. Images were collected using a 1.05 NA objective in 1× PBS (4.5-fold expansion). **c**, SRS spectral analysis of the cell area (blue) and extracellular vesicles (EVs, red) in the human lung organoid. Left, selected areas for spectral analysis representing the cell (blue) and EVs (red). Right, background-subtracted SRS spectra. Arrow indicated a side peak around 2865 cm^-1^ contributing from the CH_2_ signal of lipids in small EVs. Raw hyperspectral SRS spectra and all single-frequency SRS images are in Supplementary Fig. 6a. **d**, Hyperspectral SRS spectrum at C-H region of a single EV in (**b**) marked by a pink arrow. **e**, Spectrally unmixed C-H channels for protein and lipids in the expanded mouse brain tissue. Right, zoom into areas outlined by the white boxes. Images were collected using a 1.05 NA objective in 1× PBS (2.0-fold expansion, original ExM gel). **f**, Overlayed image of protein and lipids in the expanded mouse brain tissue. Images were collected using a 1.05 NA objective in 1/25× PBS (2.6-fold expansion, original ExM gel). **g**, Magnified area outlined by the white box in (**f**) of transverse cross-section of an individual axon. Down, corresponding plot of the intensity of proteins and lipids along the yellow box long axis. Shaded areas in (**a, c, d**) indicate the s.e.m. from different pixels. Scale bars, 500 nm in (**g**); 2 μm in (**a, b, e**); 5 μm in (**f**); 10 μm in (**c**).

EVs have been growingly recognized to play crucial roles in diverse biological systems to shuttle molecules across cells, showing promise to aid in disease diagnosis or to be engineered to deliver therapeutic payloads^38^. However, imaging EVs at the single-vesicle level remains a challenging task due to their small size and potential labeling bias^39^. Here, we have demonstrated that MAGNIFIERS enables analyzing chemical compositions of single exosomes in their spatial context without labeling (Fig. 3a-d). It is noteworthy that we only observed lipids in small EVs at submicron size, but not others such as large microvesicles and apoptotic bodies (Supplementary Fig. 6d). This could be explained by the differential lipid compositions of EVs determined by their biogenesis processes^35^. In this regard, the existence of CH_L_ signals may work as a marker to identify EV biogenesis.

We also observed interesting patterns from the CH_L_ channel in the expanded PFA-fixed mouse brain under varied expansion levels of up to 4.5-fold (Supplementary Fig. 7). For instance, by looking at the white matter in cerebellum, astounding lipid architectures were captured at the edge of myelin fibers which could be visualized by the protein channel (Supplementary Fig. 7a). Transverse cross-section of myelinated axons was clearly unveiled in the CH_L_ channel across the brain tissue (Fig. 3e-f). Axon fibers are enwrapped by myelin, a multilayered membranous sheath that serves as an electrical insulator for the axon to provide rapid conduction of action potentials. Myelin contains a high content of lipids made of large amounts of cholesterol and phospholipids (e.g. sphingomyelin), a very similar composition to the exosomes studied above. Zooming into an individual myelinated axon, lipids signal appeared as a hollow structure with a core axon and a myelin covering (Fig. 3g).

Overall, MAGNIFIERS can not only examine the ultrafine structures but also advance in revealing their chemical compositions, such as lipids and nucleic acids.

### High-resolution metabolic imaging of Huntingtin aggregates

Metabolic activities are the outcome of gene expression and relate to many dysfunctions and diseases. Bioorthogonal chemical imaging is a new paradigm which utilizes ‘chemical bonds’ as tiny vibrational tags for profiling small metabolites in cells and animals^40–42^. Different from bulky fluorophores which easily affect the function of the attached small metabolites, vibrational tags are minimally perturbative to normal biological functions. Among them, carbon-deuterium bonds are particularly effective with little perturbation, broad accessibility and residing in the bioorthogonal spectral window (where no endogenous biomolecules vibrate). Leveraging the power of small vibrational tags, MAGNIFIERS shall be able to gauge nanoscale features of metabolic activity.

Abnormal protein aggregation is associated with many neurodegenerative diseases. Aggregation-prone polyglutamine (polyQ) protein is directly linked to Huntington’s disease pathogenesis. Recent evidence suggests metabolic dysregulation in aggregates, which could be a potential target for future therapies^43^. Being able to visualize fine structures of protein synthesis in small aggregates in the cellular environment could gain mechanistic insights and uncover therapeutic opportunities. While super-resolution fluorescence imaging techniques have been extensively applied to study polyQ aggregates on resolving fine details of structures^44, 45^, they have failed to provide information on metabolic activities of these aggregates. Using deuterium-labeled amino acids (D-AAs)^41^, here we demonstrate the application of MAGNIFIERS for metabolic imaging by studying protein synthesis of polyQ. After concurrently expressing polyQ proteins and labeling newly synthesized proteins with D-AAs in HeLa cells (Supplementary Fig. 8a), we fixed the cells, then gelled and linearly expanded them ~3.4 fold in each dimension (Fig. 4a). Formation of PolyQ aggregates with different patterns (such as large inclusion bodies (IBs), fibrils and branched clusters) were observed in both the cytosol and nucleus through imaging the CH_3_ peak at 2941 cm^-1^ or the CD peak at 2135 cm^-1^ (Fig. 4b-d). The improved spatial resolution is beneficial for resolving closely packed small clusters (Fig. 4c, inset) and the heterogeneous patterns of individual protein aggregates (Fig. 4d, inset).

**Fig. 4.**
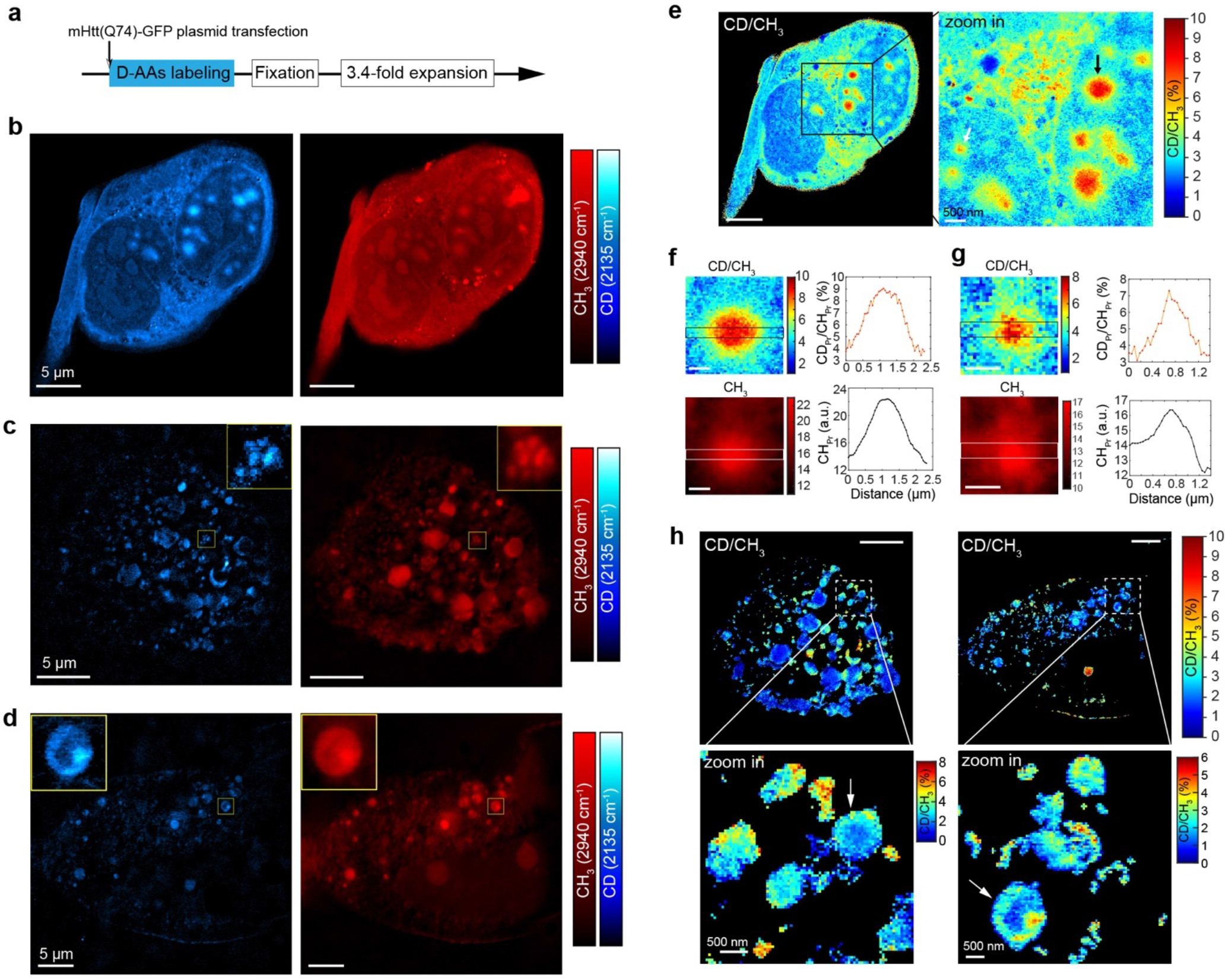
Super-resolution metabolic imaging of newly synthesized protein in Huntingtin aggregates. **a**, Deuterated amino acids (D-AAs) labeling in time with simultaneous expression of mutant huntingtin (mHtt74Q-GFP) proteins for 48 hrs. The cartoons display the experimental pipeline of plasmid transfection, medium exchanges, fixation and expansion. **b**-**d**, Representative ROIs of CH_3_ images at 2941 cm^-1^ and CD images at 2135 cm^-1^ after expansion. Up in (**c, d**), zoom into the region outlined by the yellow box. **e**, Ratio images of CD/CH_3_ for ROI in (**b**). Right, zoom into the region outlined by the black box. Arrow pointed out representative aggregates analyzed in (**f**-**g**). (f) Analysis of the aggregate pointed with black arrow in (**e**). **g**, Analysis of the aggregate pointed with white arrow in (**e**). **h**, Ratio images of CD/ CH_3_ for ROI in (**c, d**). Down, zoom into the regions outlined by the dotted box. All images were collected with a 1.05 NA objective in 5× PBS (3.44-fold expansion). Scale bars, 5 μm in (**b**-**d, e** (left), **h** (up)); 500 nm in (**f**-**g, e** (right), **h** (down)).

Interestingly, the spatial distributions of new proteins (CD channel) and old proteins (CH_3_ channel) were not always consistent. Quantitatively, the protein synthesis dynamics can be evaluated via the ratio of CD intensity over CH_3_ intensity (Fig. 4e-i). In a cell with multiple IBs located in the nucleus, we observed higher CD/CH_3_ values in the core of both large (Fig. 4f) and small (Fig. 4g) IBs. This may suggest different phases of aggregates formation: the initial seeding of aggregates rely on active loading of misfolded polyQ and later aggregates expansion recruit not only misfolded polyQ but more other surrounding normal proteins. However, in cells with abundant aggregate formation and severely impaired cellular structures, we observed higher metabolic activities close to the boundary of some our aggregates (Fig. 4h), which may be generated through fragmentation of large aggregates. In addition, through labeling different time periods of polyQ expression, we can monitor the growth trajectory of aggregates. For example, with pulse-chase labeling using regular medium first and D-AAs replaced medium later, we found large clusters of aggregates are metabolically inactive in a long time period (Supplementary Fig. 8b).

Overall, MAGNIFIERS platform can map metabolic activities of protein synthesis and turnover with sub-diffraction-limited spatial resolution of ~100 nm. We demonstrated its utility in the context of studying polyQ aggregates inside mammalian cells, which shows various nanoscale patterns of new protein synthesis. The observed features (core and boundary) of polyQ aggregates all have sub-diffraction-limited length scales. While we demonstrated with D-AAs here for studying protein synthesis, this principle can be applied to many other cases such as tracing lipid uptake with deuterium-labeled fatty acids and detecting DNA synthesis with 5-ethynyl-2’-deoxyuridine (EdU).

### Nanoscale Raman dye imaging

Multiplexed super-resolution protein imaging in 3D tissues remains a technical challenge. Limited by the broad excitation and emission bandwidth of fluorescence, it is fundamentally difficult for regular super-resolution techniques to detect more than 4 targets. While researchers have demonstrated super-resolution imaging of multiple targets using direct STORM^46^, gel embedding and expansion^26, 47^, and DNA-PAINT^48, 49^, they all relied on several rounds of staining, imaging, stripping and/or restaining. These cyclic methods could be artifact-prone, considering fine structures are vulnerable to non-linear histological changes among cycles and necessitate higher precision on image alignment. To overcome these technical difficulties, we plan to employ a one-shot multiplexing approach, by leveraging much narrower vibrational transitions.

Recently, we developed a novel multiplexing method of electronic pre-resonant SRS (epr-SRS) which detects electronically coupled vibrational modes in far-red absorbing dyes with sub-micromolar sensitivity. Probe-wise, a specially engineered MARS dye palette with π-conjugated triple bonds has been developed, which generates single and sharp peaks at the cell-silent region^6, 8^. Epr-SRS enables Raman detection of interesting biomarkers ranging from proteins, transcripts, and enzyme activities^6, 50^. In particular, immuno-eprSRS with MARS dye-conjugated antibodies has emerged as an encouraging platform for one-shot multiplexed protein imaging^6, 11^, bypassing technical troubles involved in cyclic techniques. Yet, it only works in a diffraction-limited paradigm.

We first studied the interplay between epr-SRS and MAGNIFY. Because the enzyme-free homogenization step enables post-expansion labeling (Fig. 1a), more epitopes may be revealed^51^, thus enhancing the labeling efficiency and overall signal size of immuno-eprSRS. We tested immuno-eprSRS in MAGNIFIERS using MARS2228 NHS ester conjugated secondary antibody (dye structure in Fig. 5a) to stain the expanded mouse brain section. However, we observed nonspecific staining backgrounds to varying degrees when the regular staining buffer of 1 × PBS was utilized (Fig. 5b, Supplementary Fig. 9a-b). Considering MARS dyes are positively charged and the gel matrix contains negatively charged carboxyl groups, we reasoned this non-specific staining might originate from an electrostatic effect. Hence, we changed the staining and washing buffers to a high-salt solution of 9× PBS (charge shielding effect) and supplemented with high-concentration Triton X-100 (10%) which has been reported to prompt antibody binding^52^. Effectively, non-specific staining was substantially mitigated as we observed correct staining pattern of Synaptophysin labeled with MARS-dye conjugated secondary antibody in mouse brain (Fig. 5c, Supplementary Fig. 9c).

**Fig. 5.**
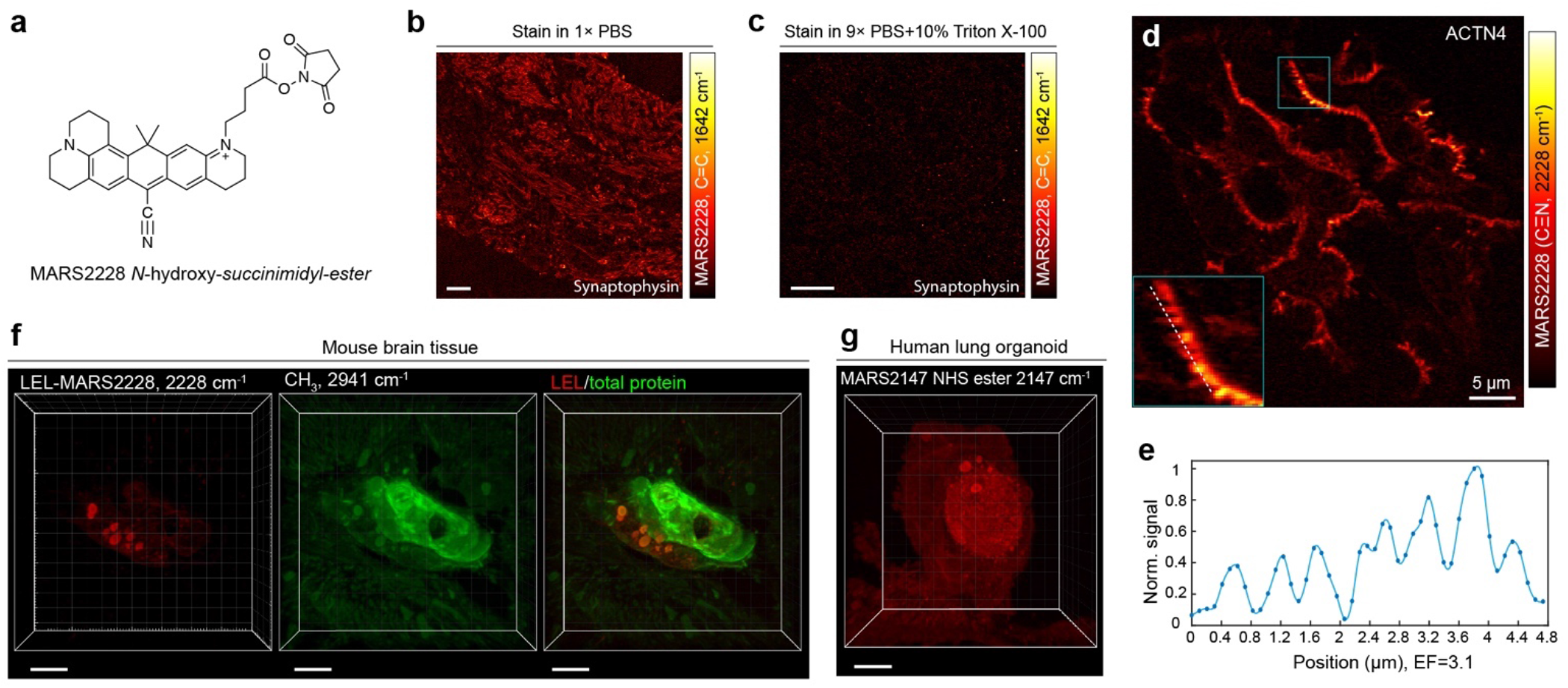
Nanoscale Raman dye imaging. **a**, Chemical structure of NHS ester functionalized MARS2228. **b**, Staining background when 1× PBS was used as the staining buffer. **c**, Correct staining patterns of Synaptophysin using 9× PBS with 10% Triton-X as the staining buffer. **d**, Immuno-eprSRS image of actinin-4 (ACTN4, specifically label tertiary podocyte foot processes) in FFPE human kidney tissue with MARS2228. Inset, zoom into the region outlined by the blue box, dotted white curve within the inset indicates the line cut analyzed below. **e**, Normalized epr-SRS signal of the nitrile mode of MARS2228 along the line cut of the inset in (**d**). **f**, 3D-rendered epr-SRS image of MARS2228-conjugated *Lycopersicon Esculentum* lectin (LEL) labeling and CH_3_ image of 2941 cm^-1^. **g**, 3D-rendered epr-SRS image of expanded human lung organoid stained with MARS2147 NHS ester dye. Images were collected using a 1.05 NA objective (**d**) in 1/50× PBS (3.1-fold expansion, original ExM gel), (**f**-**g**) in 1× PBS (4.5-fold expansion). Scale bars, 10 μm in (**b, c**); 5 μm in (**d, f, g**).

We next demonstrated the ability of MAGNIFIERS in visualizing nano-architecture of specific proteins with MARS dyes. By performing immuno-eprSRS of actinin 4 (ACTN4) in human kidney FFPE tissue with MARS dye conjugated secondary antibody, we observed ultrafine structures of the podocyte foot process by targeting the nitrile mode of MARS2228 (Fig. 5d, Supplementary Fig. 10a), consistent with standard immunofluorescence result^21^. The periodic pattern of the foot processes, which have a width around 200 nm, was clearly visualized in the line plot (Fig. 5e, Supplementary Fig. 10a). Going beyond immunolabeling, other labeling approaches such as glycoprotein labeling with lectins (Fig. 5f) and pan-proteome staining^32^ with NHS ester or maleimide functionalized MARS probes (Fig. 5g, Supplementary Fig. 10b-d) are generally viable to MAGNIFIERS in a 3D fashion (Supplementary Videos 3-4). Of note, MARS dye staining and imaging does not compromise label-free imaging of endogenous biomolecules, as shown in Fig. 5f. Collectively, these results demonstrated the applicability of Raman dye imaging with high molecular specificity in MAGNIFIERS.

### One-shot highly multiplexed nanoscale imaging

We next set out to showcase the one-shot highly-multiplexed nanoscale imaging capability of MAGNIFIERS in the mouse brain tissue section. To pursue better image contrast, we employ a simple but effective signal amplification strategy by using recurrent staining of MARS-conjugated secondary antibodies. In detail, as illustrated in Supplementary Fig. 11a, two different secondary antibodies, each of which binds to the other, will be applied alternatively after regular immunostaining process. Since postexpansion staining was utilized, the apparent size of the antibody is much smaller compared to the targeted structure in expanded samples. Therefore, recurrent staining rounds are unlikely to affect the interpretation on the shape and size of targeted proteins, as has been calibrated through standard immunofluorescence^53^. We tested this amplification method on mouse brain tissue via targeting synaptophysin, a protein found in synaptic vesicles and commonly used as a marker of pre-synaptic structures^54^. After additional 4-round recurrent staining of MARS2228-conjugated secondary antibodies, we observed substantial signal enhancement without noticeable background (Supplementary Fig. 11b-c). Quantitatively, epr-SRS signal of the skeletal C=C mode of MARS2228 increased by 1.7-fold and 2.8-fold after 2-round and 4-round amplification, respectively (Supplementary Fig. 11d).

With this strategy, we acquired three epr-SRS channels of synaptophysin, α-tubulin and postsynaptic density-95 (PSD95) each with additional 4-round staining in expanded mouse brain tissue. Fluorescence channels are also readily retained for nucleus (DAPI), microtubule associated protein 2 (MAP2) and *Lycopersicon esculentum* (tomato) lectin (LEL). Adding up the chemical imaging of CH_Pr_ and CH_L_ channels, we achieved a total of eight channels in volume in a single round (Fig. 6a), with an expansion of ~4.5 fold in each dimension (in 1× PBS, effective resolution of ~73 nm). Such a one-shot method bypasses the error-prone registration procedure and is particularly favorable for precise profiling of nanoarchitectures in thick volumes.

**Fig. 6.**
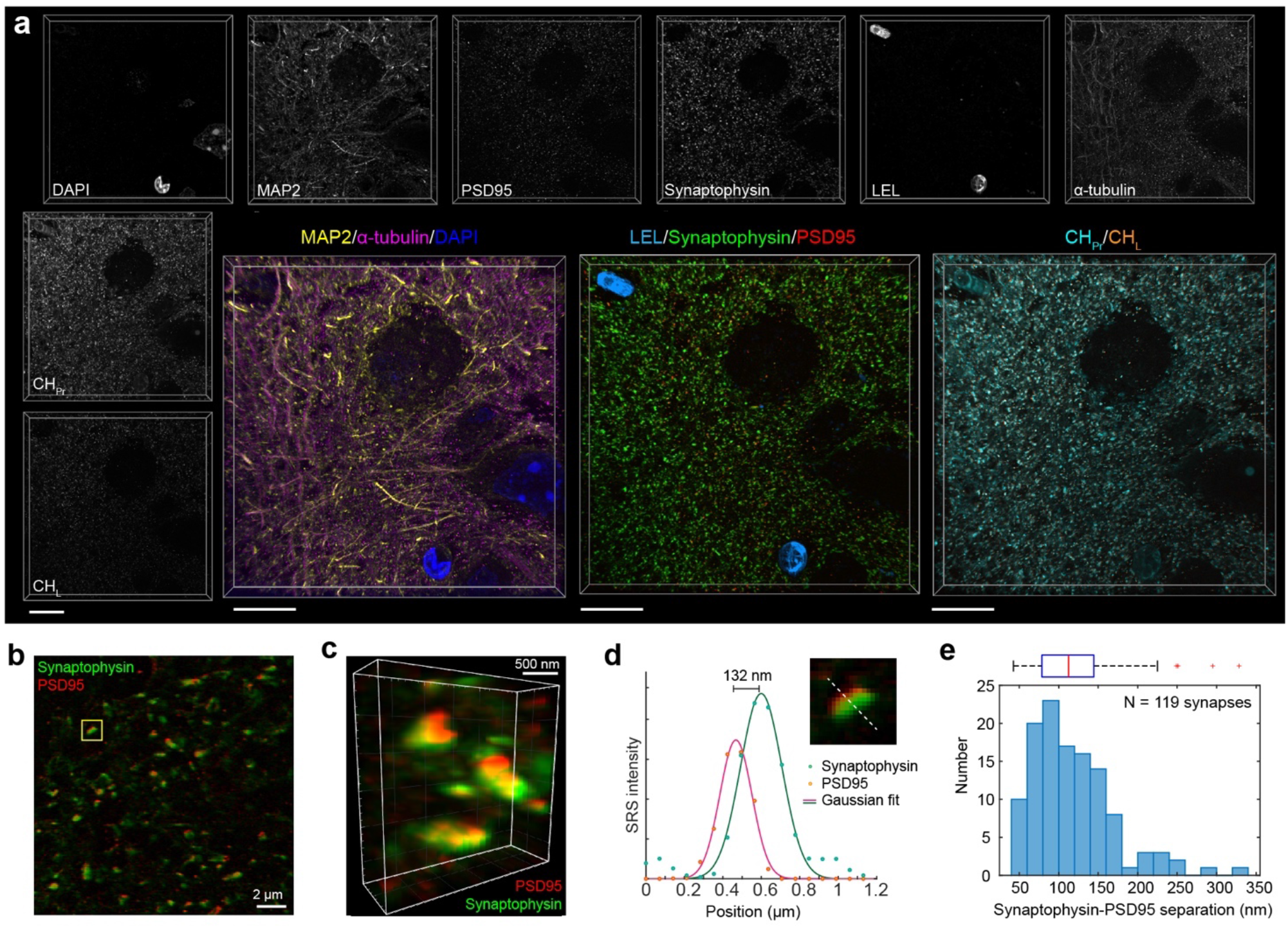
One-shot highly multiplexed nanoscale imaging. **a**, One-shot 8-plex 3D-rendered nanoscale imaging of post-expansion mouse brain slice. Fluorescence: DAPI (total DNA), microtubule associated protein 2 (MAP2, Cy3), *Lycopersicon Esculentum* lectin (LEL-DyLight 649, blood vessels). SRS: Synaptophysin (MARS2228, synapse vesicles), α-tubulin (MARS2176), postsynaptic density protein 95 (PSD95, MARS2147, post-synaptic membrane), CH_Pr_ and CH_L_. Voxel size is 0.318×0.318×1 μm in post-expansion distance. **b**, Single-plane image of overlayed synaptophysin and PSD95 channels. **c**, Zoom-in of 3D-rendered overlayed synaptophysin and PSD95 image. Yellow box outlined zoom-in image in (d) for separation distance analysis. **d**, A representative image of single synapses and corresponding plot of the staining intensity of PSD95 and synaptophysin along the dotted line. Solid lines, Gaussian fits. **e**, Statistical analysis of Synaptophysin-PSD95 separations (N=119 synapses). Above is the box plot of the separation distances. In the box plot, the center indicates the median; the bottom and top edges of the box indicate the 25^th^ and 75^th^ percentiles, respectively; the whiskers extend to the minimum and maximum data points; the outliers are plotted individually using the ‘+’ marker symbol. Images were acquired with a 25× objective in 1× PBS (4.5-fold expansion). Scale bars, 10 μm in (**a**), 2 μm in (**b**), 500 nm in (**c**).

To showcase the nanoscale spatial precision, we focused on two synaptic marker proteins of synaptophysin (marker of synaptic vesicles) and PSD95. The post-ExM image explicitly revealed synaptophysin juxtaposed with PSD95 puncta (Fig. 6b-c, Supplementary Video 5). The separation distance between synaptophysin-PSD95 pairs were measured after the Gaussian fitting of the signal intensity along the line perpendicular to the synaptic cleft (Fig. 6d). The synaptophysin-PSD95 separation was measured as 118±52 nm (Fig. 6e, mean±s.d., n=119 synapses) in MAGNIFIERS. Our result is close to the prior measurement of separations between postsynaptic densities and synaptic vesicles of around 150 nm using super-resolution fluorescence imaging^54^. To our best knowledge, vibrational imaging has not been applied to visualize individual synaptic features before. Overall, we expect MAGNIFIERS will provide a parallel and scalable way to interrogate nano-architectures in 3D spatial context with molecular specificity.

## Discussion

SRS microscopy and ExM, two emerging techniques, have been synergistically integrated in MAGNIFIERS, providing niches that are complementary to existing SRS or ExM imaging alone. On one side, MAGNIFIERS offers a simple and general way to achieve super-resolution Raman imaging; on the other side, it complements the prevalent fluorescence-based nanoscopy. Leveraging the wide-ranging capabilities of SRS microscopy, we have demonstrated three classes of applications: label-free chemical imaging, metabolic imaging, and multiplexed imaging with nanoscale precision (Fig. 1a). For chemical imaging (Figs. 2-3), MAGNIFIERS provides label-free access to visualize proteins, lipids and DNA biomolecules with effective resolution up to ~41 nm. We have demonstrated this utility in varied biologically meaningful scenarios, such as visualization of the ultrastructure of basal bodies and investigation of the chemical composition of EVs in human lung organoids. We foresee that MAGNIFIERS will contribute single-vesicle level understanding of EVs in vivo. For metabolic imaging (Fig. 4), we studied protein synthesis and turnover on protein aggregates using deuterated amino acid labeling. In multiplexed imaging with Raman dyes (Figs. 5-6), MAGNIFIERS offers a one-shot optical approach to simultaneously map multiple targets in 3D in mouse brain sections. Finally, these applications are additive. For instance, chemical imaging is compatible with imaging of metabolic activities or postexpansion Raman dye imaging, providing a comprehensive view of the chemical content of the sample.

A recent technique called VISTA combines label-free SRS imaging of protein with a special ExM protocol named magnified analysis of the proteome (MAP)^29, 55^. Compared to VISTA, MAGNIFIERS demonstrates the full potential of expansion SRS microscopy on a much broader range of biological specimens by leveraging the most recent advancement in ExM^30^. Foremost, unlike VISTA which requires the special acrylamide co-fixation and only reportedly works on mouse brain tissues^26, 29^, MAGNIFIERS is not limited to soft tissue such as the brain and features general sample compatibility across a broad range of tissue types and fixation methods. We have shown that mechanically tough tissues such as FFPE human kidney can be robustly expanded and imaged by MAGNIFIERS, thus making it readily applicable to the vast archive of human specimens. Moreover, by virtue of more effective preservation of biomolecules (near 100%) and tunable expansion level, our approach offers better detection sensitivity as compared to VISTA. This not only enables nanoscale label-free imaging of protein, nucleic acid, and lipids, but also allows us to open up nanoscale metabolic imaging and highly multiplexed Raman dye imaging, which generally have weaker signals than label-free applications. Lastly, MAGNIFIERS provides finer spatial resolution over VISTA by 1.6-fold through leveraging excellent reversible linear expansion of ~7.2-fold. In contrast, VISTA only allowed up to ~4.5-fold expansion. Altogether, MAGNIFIERS is currently the only technique that truly holds the value behind the concept of Expansion SRS microscopy: (1) true label-free nanoscale imaging of protein, nucleic acid, and lipids; (2) superresolution vibrational metabolic imaging; and (3) super-multiplexing capability. In contrast, VISTA only demonstrated part of (1) but none of (2) and (3).

Future improvements and opportunities reside in several fronts. First, the full expansion potential of the MAGNIFY gel has not been thoroughly utilized. To pursue higher effective resolution, femtosecond pulse excitation could be a direct way to enhance broadband SRS signals such as C-H and C-D bonds by an order of magnitude^56^. In this way, full expansion of ~10-fold in water will be viable for label-free protein imaging. Moreover, for Raman dye imaging, other DNA-based signal amplification techniques such as immuno-SABER^47^ and isHCR^57^ are expected to boost the signal more efficiently. Particularly, these amplification strategies will further increase the multiplexity with a practical goal of over 20 channels^7^. Second, specimens become optically transparent after expansion and thus MAGNIFIERS has promising applications for thick specimens of millimeter scales, considering successful precedents in integrating tissue clearing with label-free SRS^58^ or highly-multiplexed Raman dye imaging^11^. This would open exciting applications to probe interactions between ultrastructure with high content over a long distance, such as with neuronal tracing. Finally, while SRS may have low throughput for profiling large-volume specimens at high precision, the reversible and size-controllable features of the MAGNIFY gel provide good flexibility to combine large volume profiling and small region-of-interest interrogation in the same sample.

## Acknowledgments

We kindly thank Lingyan Shi for the assistance in animal experiments, thank Chenyi Mao and Gianluca Oyarzún for helpful discussions. W.M. acknowledges support from NIH (R01 GM128214, R01 GM132860 and R01 EB029523). Y.Z. acknowledges support from Carnegie Mellon University, DSF charitable foundations, U.S. Department of Defense DoD VR190139, NIH Director’s New Innovator Award DP2 OD025926-01. X. R. acknowledges support from DSF charitable foundations and the Department of Biomedical Engineering at Carnegie Mellon University. P.W. acknowledges T32 pre-doctoral training grant (Biomechanics in Regenerative Medicine, BiRM) from the National Institute of Biomedical Imaging and Bioengineering of NIH.

## Author contributions

LS performed SRS imaging experiments and analyzed related data. AK, BRG, FF and YZ performed all expansion experiments and related characterizations. ZC contributed to image processing of multiplex SRS images. PW and XR prepared lung organoid samples. YM synthesized MARS probes. LS, YZ and WM conceived the concept and wrote the manuscript with input from all authors. YZ and WM supervised the project.

## Data and materials availability

All data supporting this work are available in the main text or the supplementary materials. All raw data are available from the corresponding authors upon request.

