## Supplementary Information for "Super-resolution vibrational imaging using expansion stimulated Raman scattering microscopy"

**This file includes:**

Methods
Supplementary Figures 1-11
Supplementary Tables 1-3
Supplementary Videos 1-5
References

### Methods

**Materials and human samples.** Detailed information regarding all reagents and equipment was described in Supplementary Table 1. MARS dyes and their derivatives (NHS-ester- and maleimide-functionalized) were synthesized as previously reported[1]. The human pathology specimens were purchased from US Biomax catalog numbers HuFPT072 (normal human kidney cortex), HuFPT015 (normal human hippocampus), HuFPT082 (normal human spleen).

**Mouse sample preparation.** Animal experimental protocol (AC-AABD1552) was approved by the Institutional Animal Care and Use Committee (IACUC) at Columbia University. All experiments using mice were conducted in strict adherence to the ethical regulations of Columbia University IACUC. Wild type male and female mice (C57BL/6, ~5 weeks old, Jackson Lab) were fully anesthetized using isoflurane, then sacrificed with cervical displacement and immediately perfused with 4% paraformaldehyde (PFA) in PBS transcranially. The brain, kidney, liver and pancreas were extracted and fixed in 4% PFA in PBS at 4 °C for 24 h. After that, the collected organs were immersed in PBS at 4 °C for 24 h to remove PFA. The organ was embedded in 7% agarose gel and sectioned into 40 µm thick coronal slices using a vibratome (VT1000S, Leica). Agarose was removed by a tweezer after sectioning.

**Culture of airway basal cells.** The normal human bronchial epithelial (NHBE) cultured in 804G-conditioned medium coated culture vessels in bronchial epithelial cell growth medium (BEGM) supplemented with 1uM A8301, 5uM Y27632, 0.2uM of DMH-1, and 0.5uM of CHIR99021[2] at 37°C with 5% $CO_2$.

**Differentiation of airway basal cells into airway organoids.** A 96-well tissue culture plate was coated with 40% (vol/vol) growth factor reduced (GFR) Matrigel in PneumaCult™-ALI Maintenance Medium. The NHBEs were resuspended in 40% (vol/vol) GFR Matrigel in PneumaCult™-ALI Maintenance Medium and added to the coated wells. 100µL PneumaCult™-ALI Maintenance Medium was placed in the wells and changed every other day. The cultures were maintained at 37°C with 5% $CO_2$ for 21 days.

**Deuterated amino acids labeling to study huntingtin protein aggregation.** HeLa cells (ATCC CCL-2) were cultured in DMEM (11965, ThermoFisher) supplemented with 10% FBS (10082, ThermoFisher) and 1% penicillin/streptomycin (1514, ThermoFisher). Cells were first seeded onto a clean coverslip and cultured with their usual culture media for 24 h, followed by transfection with 200 ng Htt-Q74 (Addgene, #40262, tagged with EGFP) plasmid using Lipofectamine™ 3000 transfection reagent in regular DMEM or CD-DMEM (see previous report[3] for recipe). After 48 hr incubation, cells were washed with DPBS and fixed with 4% PFA for 15 min. Cells were then expanded following the protocol described above.

**Protein-MARS probe conjugation.** NHS-ester-functionalized MARS probes were stored at a concentration of 10 mM in DMSO under -20 °C, protecting from light and moisture. To perform protein-dye conjugation, dye solutions were first diluted in DMSO to a concentration of 2 mg/mL. Conjugation buffer was prepared as 0.1 M $NaHCO_3$ in PBS buffer with pH adjusted to 8.3. Highly cross-adsorbed secondary antibodies were buffer exchanged and concentrated to 2 mg/mL in the conjugation buffer. A 50 µL dye-NHS solution was slowly added to a 0.5 mL secondary antibody solution under stirring. For primary antibody labeling, the protein concentration was adjusted to 1 mg/ml and the molar ratio of dye/protein was usually 10-15. Lectins were first dissolved in the conjugation buffer as 2 mg/ml. For *Lycopersicon Esculentum* Lectin labeling, a 25 µL 2 mg/ml dye-NHS solution was added to a 0.5 mL 2 mg/ml LEL solution. Reactions were all incubated at room temperature for 1 h under constant mild stirring. Labeled proteins were further separated from unreacted dyes by gel permeation chromatography using Sephadex™ G-25 (G25150 SIGMA) resins with a column of 1-cm diameter and over 12-cm length. Purified protein solution was centrifuged to remove potential precipitates and further concentrated with Amicon® Ultra Centrifugal Filters (UFC501096, EMD, Millipore). A final concentration of ~2 mg/mL protein solution (for secondary antibodies and lectins) were prepared in stocking buffer (30% glycerol and 5 mM sodium azide in PBS) and stored at -20 °C. The degree of labeling (DOL, i.e. dye-to-protein ratio) were measured with UV-Vis spectrum using Tecan Infinite 200 Reader with a NanoQuant Plate. DOL on secondary antibodies is about 3.

**Stimulated Raman scattering (SRS) and fluorescence integrated imaging platform.** SRS and fluorescence (both confocal and two-photon) imaging were performed on an inverted laser scanning microscope (Olympus FV1200) using a 25× water-immersion objective lens (Olympus XLPlan N, 1.05 NA, MP, WD = 2 mm) or a 60× IR-coated water-immersion objective lens (Olympus UPlanApo/IR, 1.2 NA).

For SRS imaging, two synchronized 6-ps lasers (called pump and Stokes beams) with 80-MHz repetition rate are provided by a picoEmerald system from APE (Applied Physics & Electronic, Inc.). Pump beam is tunable from 720-990 nm through both temperature control of the nonlinear crystal and a Lyot filter. Stokes beam is fixed at 1064.2 nm. The intensity of the Stokes beam was modulated sinusoidally by a built-in electro-optic modulator (EOM) at 8 MHz with a modulation depth of more than 90%. Spatially and temporally-overlapped pump and Stokes beams were coupled into the laser-scanning microscope. After passing through the specimens, forward-going pump and Stokes beams were collected with an IR-coated oil condenser (1.4 NA, Olympus). Stokes beam were completely filtered with two high-optical-density bandpass filter (890/220 CARS, Chroma Technology) and transmitted pump beam was detected by a large-area (10 mm×10 mm) Si photodiode (FDS1010, Thorlabs). The output current of the photodiode was then sent to a fast lock-in amplifier (HF2LI, Zurich Instruments) for signal demodulation. For C-H and C-D bonds, the laser power was set as $P_{\text{pump}}$=67-100 mW, $P_{\text{Stokes}}$=100-150 mW, and the pixel dwell time is 60-80 μs and the corresponding time constant of the lock-in amplifier is 30-40 μs. For eprSRS imaging of MARS dyes, the laser power was set as $P_{\text{pump}}$=17 mW, $P_{\text{Stokes}}$=67 mW; and images were generated through Kalman filtering of 10-30 serial frames with the pixel dwell time of 4 μs, and the time constants of lock-in amplifier were chosen as 2-4 μs. For volumetric imaging, the step size in z was 1-2 μm.

For two-photon fluorescence, DAPI dye was excited by the SRS pump laser at 760 nm. The backward fluorescence was detected after passing through a 690-nm short-pass filter, reflected by a 570-nm long-pass dichroic with a collection band of 410-490 nm. For confocal fluorescence, green channel is excited by argon laser (488 nm) with a collection band of 505-520 nm; red channel is excited by HeNe(G) laser (543 nm) with a collection band of 560-620 nm; far-red channel is excited by LD laser (635 nm) with a collection band of 655-755 nm. Multichannel photomultiplier tube (PMT) was used for fluorescence detection. Pixel dwell time was set as 2-4 μs.

**Tissue section recovery and heat treatment**. Formalin-fixed paraffin-embedded (FFPE) clinical samples were washed in the following solutions 2 times for 3 minutes each at room temperature (RT): xylene, 100% ethanol, 95% ethanol, 70% ethanol, 50% ethanol, and doubly deionized water. For samples that were stained prior to gelation, tissue slides were placed in 20 mM sodium citrate solution (pH 8) at 100 ºC. The container was transferred to a 60 ºC container for 30 minutes.

**Permeabilization of fixed tissues.** PFA fixed tissues (mouse brain, liver, kidney; HeLa cells; and lung organoid) were permeabilized for 1 hour with 1% C12E10/1xPBS or 1%PBST at RT prior to gelation with the MAGNIFY protocol.

**Protein anchoring with ProExM.** A stock solution of Acryloyl-X, s.e.m. (6-((acryloyl)amino)hexanoic acid, succinimidyl ester, AcX) was prepared by dissolving in anhydrous DMSO to a concentration of 10 mg/mL. The solution was then aliquoted and stored in a desiccated environment at −20 °C. Tissue slides were incubated with 0.05 mg/ml AcX diluted in 1x PBS buffer overnight at 4°C.

***In situ* polymer synthesis of samples with ProExM.** A gelling solution based on a modified version of the expansion pathology (ExPath) protocol was used containing 15% (w/v) SA, 5% (w/v) AA, 0.1% (w/v) Bis, 11.7% (w/v) NaCl, and 1× PBS was prepared in water. The chemicals 4HT, TEMED, and APS were added to the gel monomer solution to a final concentration of 0.01% (w/v) 4HT, 0.2% (v/v) TEMED, and 0.2% (w/v) APS. After mixing the solution was vortexed, and tissues were incubated with the gelling solution for 30 minutes at 4 ºC to allow the monomer solution to diffuse into the tissue while preventing premature gelation. A gelling chamber was then constructed around the tissue, consisting of spacers cut from #1.5 cover glass and a glass slide on top. The samples were incubated overnight in a humidified container at 37 ºC to complete gelation.

***In situ* polymer synthesis of samples with MAGNIFY.** A monomer solution made of 4% DMAA (v/v), 34% SA (w/v), 10% AA (w/v), 0.01% Bis (w/v), 1% NaCl (w/v), and 1x PBS or the modified ExPath monomer solution, or a monomer solution made of 8% DMAA (v/v), 30% SA (w/v), 5% AA (w/v), 0.01% Bis (w/v), 1% NaCl (w/v), and 1x PBS was prepared and stored at 4C prior to synthesis. Prior to gelation, the chemicals 4HT, APS, TEMED, and methacrolein were added to a final concentration of 0.2-0.25% (w/v) APS, 0-0.25% (v/v) TEMED, 0.001% 4HT (w/v), and 0.1% (v/v) methacrolein immediately prior to gelation. After mixing the solution was vortexed, and tissues were incubated with the gelling solution for 30 minutes at 4 ºC to allow the monomer solution to diffuse into the tissue while preventing premature gelation. A gelling chamber was then constructed around the tissue, consisting of spacers cut from #1.5 cover glass and a glass slide on top. The samples were incubated overnight in a humidified container at 37 ºC to complete gelation.

**Protease digestion of samples.** After gelation, blank gel surrounding the tissue was trimmed from the samples. Samples were then incubated in the ExPath homogenization buffer (50 mM Tris (pH = 8), 25 mM EDTA, 0.5% w/v Triton X-100, 0.8M NaCl) with Proteinase K diluted by 1:400 (final concentration 2 units/mL). Mouse brain samples were then homogenized at RT for 1-2h. Homogenized samples were then washed 3 times with 1x PBS at RT.

**Sample digestion and expansion with MAGNIFY.** After gelation, blank gel surrounding the tissue was trimmed from the samples and tissue was cut into smaller pieces. Samples were then incubated in the homogenization buffer (1-10% w/v SDS, 8M Urea, 25 mM EDTA, 2× PBS, pH 7.5 at RT) for1-48h at 80-95ºC with shaking. Homogenized samples were then washed 3 times with 1x PBS at RT, followed by at least 3 washes in 1% C12E10/1xPBS or 1% PBST at RT or 60 ºC to remove remaining SDS.

**Estimation of expansion factor.** Expansion factors estimated using average nuclear surface area or by matching features in pre- versus post-expansion images. For average nuclear surface area estimates, Images of DAPI stained samples were obtained using a CFI Plan Apo Lambda 10× (NA 0.45) prior to gelation and after homogenization and expansion. Nuclear surface areas were determined using FIJI/ImageJ. To calculate the linear expansion factor, the square root of the ratio of the average post-expansion nuclear surface area to average pre-expansion surface area was calculated. For specimens with pre-expansion images, immunostained samples were imaged using a CFI Plan Apo Lambda 4× (NA 0.2) and CFI Plan Apo Lambda 10× (NA 0.45) objective. After gelling and homogenization, expanded tissue pieces were imaged using a CFI Plan Apo Lambda 4× (NA 0.2) and a CFI Plan Apo Lambda 10× (NA 0.45) objective. Regions of interest in post-expanded images were matched to pre-expansion regions of interest and the measurement tool in NIS elements or FIJI/ImageJ was used to measure features sizes in both pre- and post-expansion images.

**Measurement of SRS resolution by imaging beads.** For the 25× objective (NA=1.05), 120 nm polystyrene Latex beads were first diluted in deionized water by 1:100 with vertexing and sonication. The diluted beads were added to 2.5% agarose at 95 °C on a heat plate. The mixed solution was quickly sealed between a glass slide and a coverslip before gel formation. For the 60× objective (NA=1.2), 100 nm polystyrene Latex beads were resuspended in deionized water by a 1:1000 dilution ratio with vertexing and sonication. Then the beads were sandwiched between a glass slide and a coverslip for one day to settle down. The beads were imaged through probing the 3056 $cm^{-1}$ band under each objective.

**$CH_L$/$CH_{Pr}$/$CH_{DNA}$ unmixing.** A previously reported spectral linear combination algorithm was used to unmix lipids, proteins and DNA signals on the C-H region. Briefly, SRS signature peaks at 2848 $cm^{-1}$, 2941 $cm^{-1}$ and 2967 $cm^{-1}$ were acquired, which bear the Raman features of C–H bonds in lipids, proteins and DNA, respectively. Following equations used to calculate unmixed $CH_L$, $CH_{Pr}$ and $CH_{DNA}$ signals. $I_{2848}$, $I_{2941}$ and $I_{2967}$ are SRS microscopic signal intensities at 2848, 2941 and 2967 $cm^{-1}$, respectively.

$$\begin{pmatrix} CH_{DNA} \\ CH_{Pr} \\ CH_L \end{pmatrix} = \begin{pmatrix} 1.29 & -0.58 & -0.007 \\ -0.66 & 1.38 & -0.105 \\ 0.14 & -0.93 & 1.09 \end{pmatrix} \begin{pmatrix} I_{2967} \\ I_{2941} \\ I_{2848} \end{pmatrix}$$

**Post-expansion immunostaining of FFPE samples and expansion.** After homogenization and washing, samples were stained with respective primary antibodies diluted to approximately 1 μg/mL in the staining buffer (9× PBS/10% Triton X-100/10mg/L heparin) overnight at room temperature. Samples were then washed 3 times with the washing buffer (1× PBS/0.1% TritonX-100) at room temperature for at least 10 minutes. Samples were then incubated in staining buffer with the relevant secondary antibodies diluted to 1-10 μg/mL overnight at RT. Samples were then washed at least 3 times with staining buffer for at least 30 minutes. After staining, samples were washed in 1× PBS for at least 30 minutes. This was repeated until the sample was fully expanded, at least three exchanges of final imaging solution.

**Brain tissue multicolor imaging.**

*Staining and imaging.* Post-expanded brain samples were stained in 2 ml Eppendorf tubes all at room temperature. Samples were incubated with primary antibodies (rabbit anti-synaptophysin, goat PSD95, mouse a-tubulin, chicken anti-TH, guinea pig anti-MAP2) in 9x PBS/10% TritonX/10mg/L heparin overnight at room temperature, followed by washing at room temperature for 10 min in excess volumes of 1xPBS/0.1% TritonX-100 three times. Samples were then stained with MARS conjugated secondary antibodies for four rounds at a concentration of approximately 10 μg/mL each in the staining buffer (5x PBS/10% TritonX) for 18-24 hrs. In the odd rounds, MARS2228-conjugated donkey anti-rabbit IgG antibody, MARS2176-conjugated rat anti-mouse IgG antibody, MARS2145-conjugated bovine anti-goat IgG were applied. In the even rounds, MARS2228-conjugated rabbit anti-donkey IgG, MARS2176-conjugated mouse anti-rat IgG, MARS2145-conjugated goat anti-bovine IgG were applied. Between rounds, samples were washed for 1 hr in excess volumes of washing buffer (5x PBS/10% TritonX) three times. After that, samples were stained with Cy3-conjugated donkey anti-chicken IgY, Alexa Fluro488 donkey anti-guinea pig IgG, Dylight649-conjugated *Lycopersicon Esculentum* lectin in 5x PBS. Gels were then washed and expanded in excess volumes of 1x PBS for 1 hr. This step was repeated 3-5 times until complete expansion. Lastly, gels were stained with DAPI in 1x PBS for 1 hr. For SRS imaging, gels were sandwiched in imaging gel (1xPBS/1.5% agarose (w/v)/ 1% propyl gallate (w/v)) between the cover slip and the glass slide.

*Data processing*. A linear-combination algorithm was applied on the raw volumetric epr-SRS data set to remove potential cross-talks between different channels[4]. Measured signals ($\boldsymbol{S}$) can be expressed as $\boldsymbol{S}=M\boldsymbol{C}$, where $\boldsymbol{C}$ is the MARS probe concentrations and $M$ is a $3\times3$ matrix determined by Raman cross-sections of MARS probes. MARS probe concentrations were therefore determined using $\boldsymbol{C} = M^{-1}\boldsymbol{S}$. All the images were volume-rendered in the mode of maximal intensity projection using Imaris Viewer (Bitplane).

*Synapse quantification.* Data of Fig. 6a was used for analysis. Synapses were identified by visual inspection. Candidate instances of closely apposed synaptophysin-stained and PSD-95 stained spots/ribbon were selected with following criteria: synapses were oriented perpendicular to the y-axis; complex assemblies of synapses (e.g., with multiple pre- or post-synaptic terminals) were rejected. For each synapse selected for inclusion in the analysis, an averaged (over a width of up to 8 pixels (~500 nm)) line profile of SRS signal was generated along the y-axis. The resulting intensity distributions were fitted to Gaussian distributions with Matlab 'fit' function. Any synapses with a resulting goodness of fit (i.e. rmsd), for either synaptophysin or PSD95, of less than 0.9 were rejected. The separation was calculated as the distance between the fitted centers of the Gaussian distributions for each synapse.

**Statistics.**

Statistical analysis was carried out using GraphPad Prism 7. Data are presented as mean±s.d. with statistical significance if required (not significant $P\geq0.05$, $*P<0.05$, $**P<0.01$, $***P<0.001$, $****P<0.0001$). All values of *n* are provided in the figure legends.

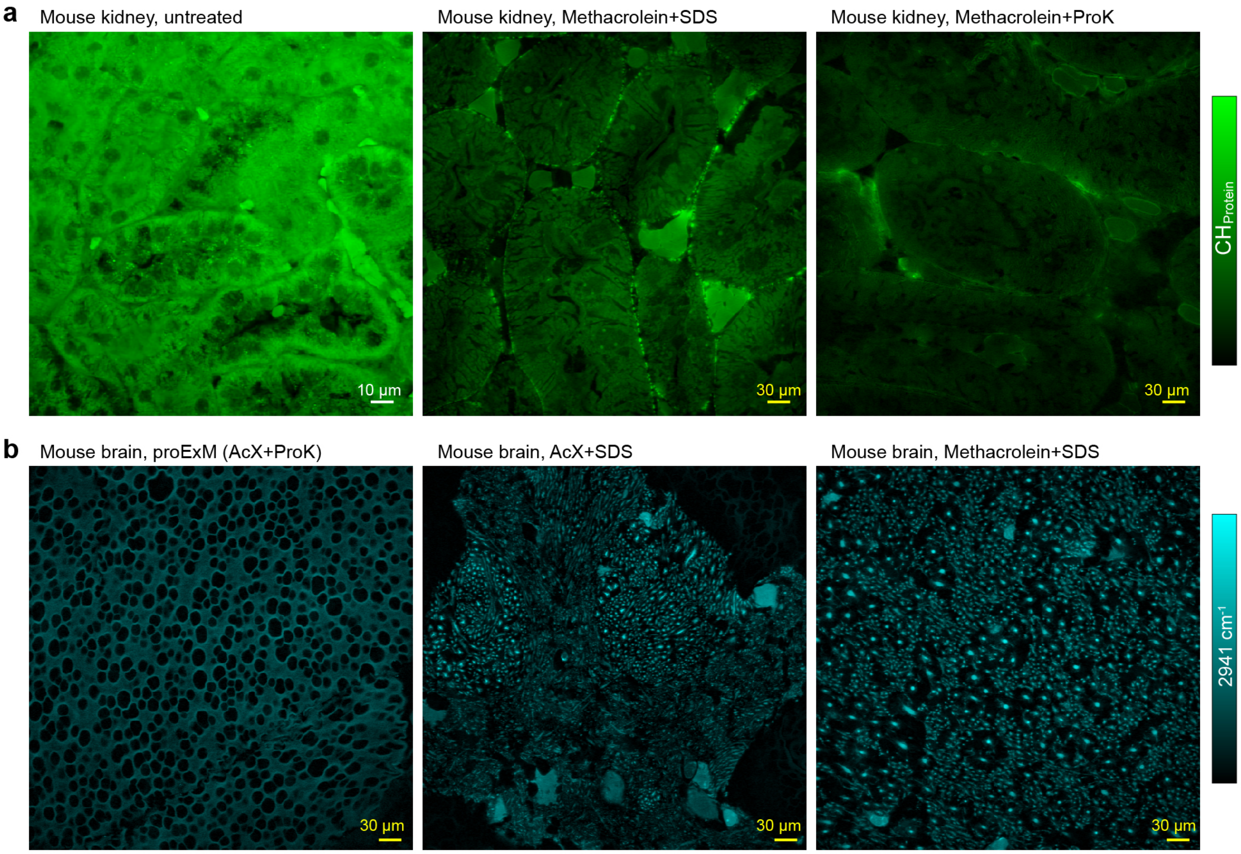


**Supplementary Figure 1. Comparison of protein retention on proExM protocol with our protocol.** **a**, SRS images of $CH_{Pr}$ of the tubules in the mouse kidney tissues. (left) untreated, (middle) expanded with methacrolein-linking and SDS/urea homogenization, (right) expanded with methacrolein-linking and Proteinase K digestion. **b**, SRS images of $CH_3$ at 2941 $cm^{-1}$ in the mouse brain tissues. (left) expanded in proExM protocol with AcX as the crosslinker and Proteinase K digestion, (middle) expanded with AcX as the crosslinker and SDS/urea homogenization, (right) expanded with methacrolein-linking and SDS/urea homogenization.

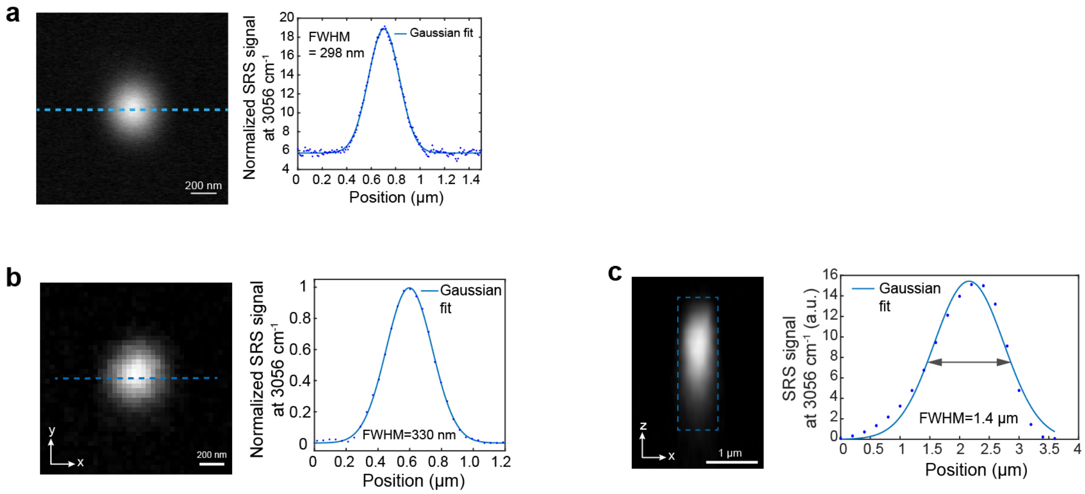


**Supplementary Figure 2. Bead quantification of the spatial resolution of SRS microscopy. a,** Calibration of lateral resolution of SRS under a 1.2 NA objective with a 100-nm polystyrene bead by measuring the C-H peak at 3056 $cm^{-1}$. Quantification of (**b**) lateral (after deconvolution) and (**c**) axial resolution of SRS under a 1.05 NA objective with a 120-nm polystyrene bead by measuring the C-H peak at 3056 $cm^{-1}$. Right graphs are cross-section profiles fitted by the Gaussian function. Scale bars, 200 nm in (**a**, **b**); 1 μm in (**c**);

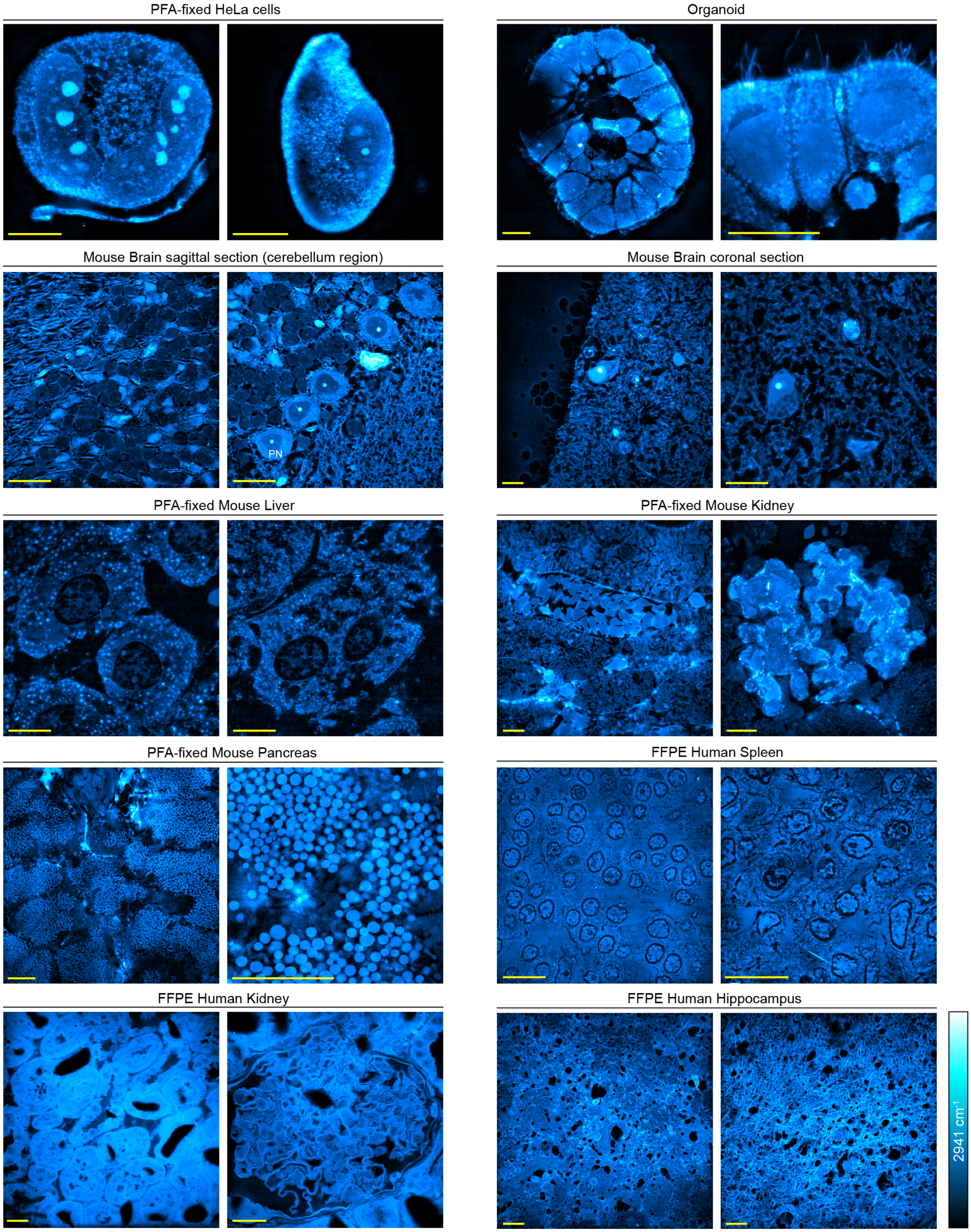


**Supplementary Figure 3. MAGNIFIERS has good sample generality.** SRS images of $CH_3$ peak at 2941 $cm^{-1}$ of PFA-fixed HeLa cells, organoids, mouse brain, mouse liver, mouse kidney, mouse pancreas, FFPE human kidney, FFPE human spleen and FFPE human hippocampus tissues. Scale bars (post-expansion), 50 μm.

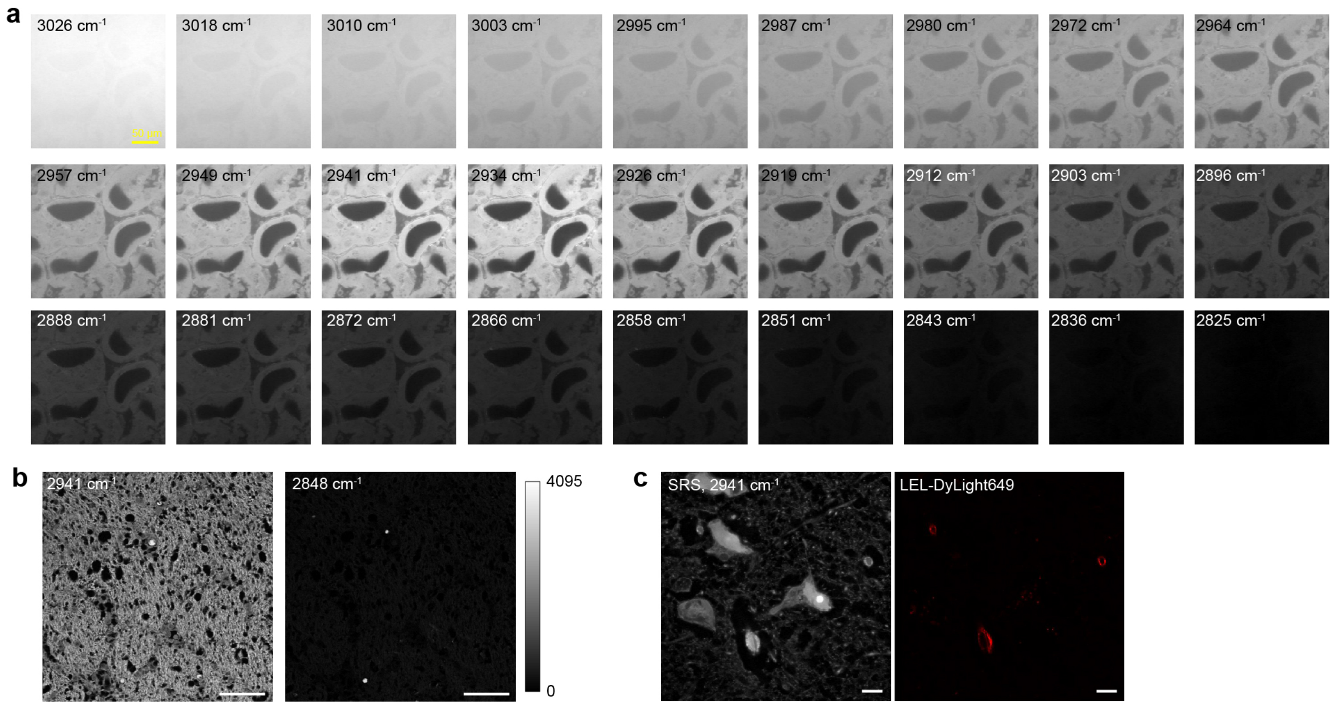


**Supplementary Figure 4. Studies on label-free protein imaging in MAGNIFIERS. a**, Images of hyperspectral SRS acquisition on the human kidney sample. The wavenumbers were labeled at the up-left corner of each image. **b**, SRS images of 2941 $cm^{-1}$ (left) and 2848 $cm^{-1}$ (right) on deparaffinized FFPE human hippocampus tissue. **c**, Expanded mouse brain tissue labeled with *Lycopersicon Esculentum* lectin (LEL)-DyLight 649. Left, SRS image of 2941 $cm^{-1}$; Right, fluorescence image of blood vessels. Expanded mouse brain in (**c**, **e**) were imaged with a 25× objective in 1× PBS with 4.5-fold expansion. Scale bars, 5 µm (post-expansion) in (**a**); 50 µm in (**b**); 5 µm in (**c**, **d**).

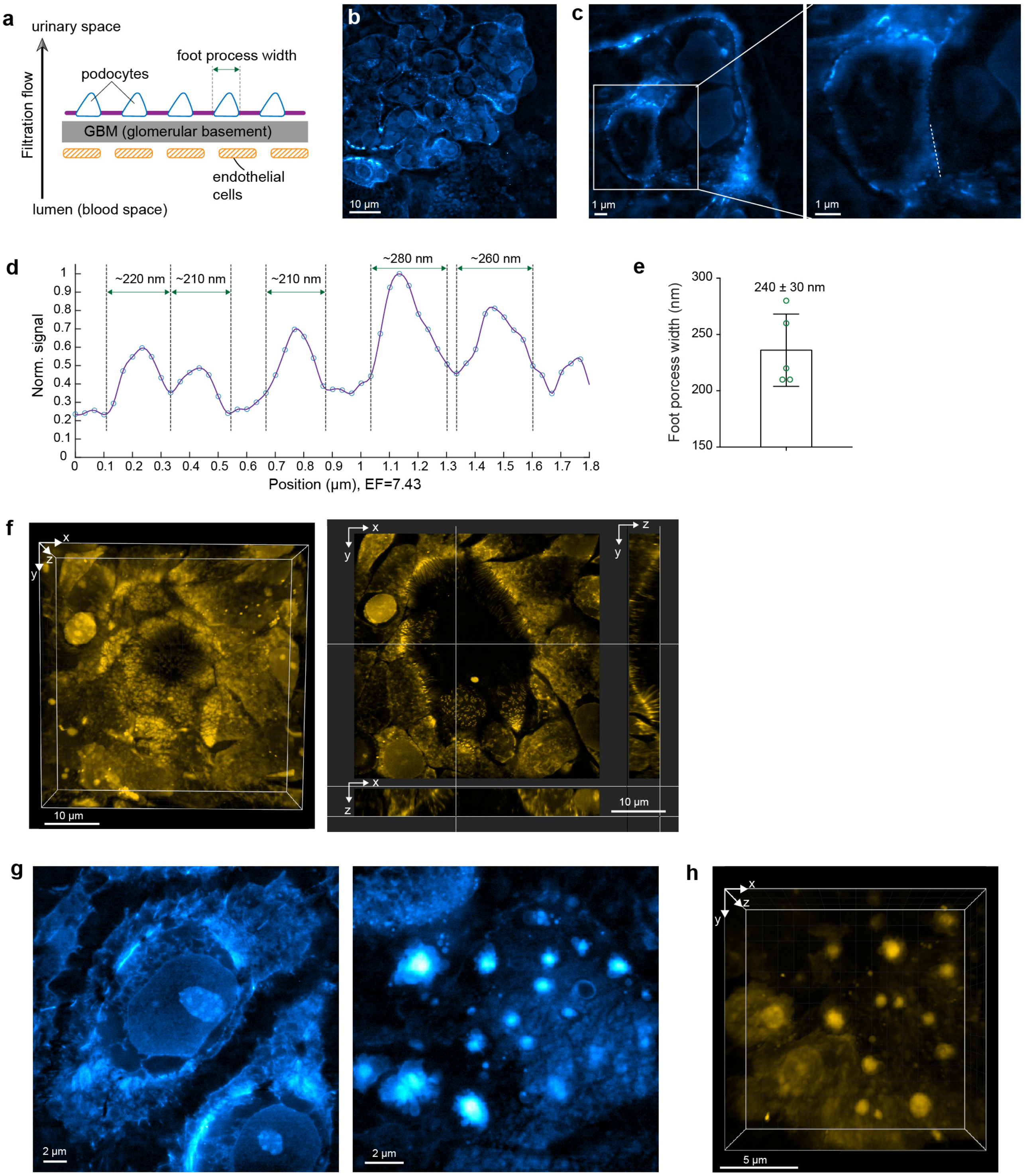


**Supplementary Figure 5. $CH_3$ image of protein reveals ultrafine structures in mouse kidney and human lung organoids. a**, Schematics of the structure of three-layered glomerular filtration barrier (GFB). GFB is an intricate structure in kidney glomeruli that builds the selective barrier between bloodstream and the urine. GFB is composed of unique epithelial cells named podocytes which extend many interdigitated foot processes in the glomeruli. **b-c**. $CH_3$ image at 2941 $cm^{-1}$ visualized ultrafine structures of tertiary podocyte foot processes in kidney glomerulus in 1/25× PBS (expansion factor, 7.43). **c**, Right, magnified image of the region outlined by the white box. **d**, Line profile of the dotted line within the right image in (**c**). **e**, Bar plot (mean±s.d.) on foot process width as marked in graph (**d**).

**f-h**, SRS images at 2941 $cm^{-1}$ of expanded human bronchial epithelial cells-derived lung organoids in 1×PBS (expansion factor, 4.5). All results were acquired with a 1.05 NA objective. Scale bars, 10 μm in (**b**, **f**); 1 μm in (**c**); 2 μm in (**g**); 5 μm in (**h**).

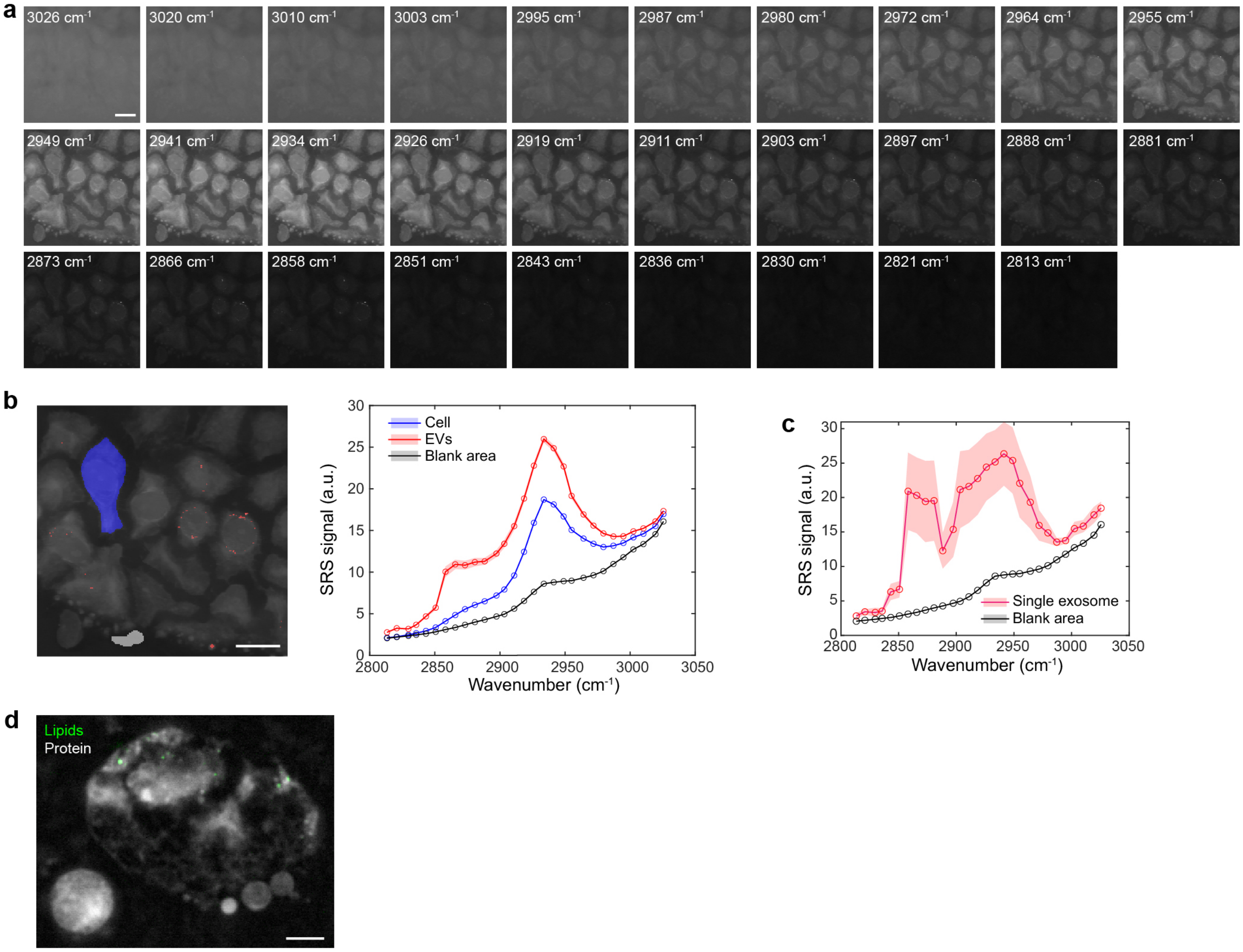


**Supplementary Figure 6. Chemical composition study in expanded human bronchial epithelial cells-derived lung organoid. a**, Images of hyperspectral SRS acquisition on the human lung organoid. The wavenumbers were labeled at the up-left corner of each image. **b**, Hyperspectral SRS spectra of cell area (blue), extracellular vesicles (EVs, red) and blank gel area (gray) of the data cube in (**a**). Selected areas were labeled on the left image. **c**, Hyperspectral SRS spectra of a single EV marked by a pink arrow in Fig. 3b and blank gel area (gray). Shaded area indicates the s.e.m. of SRS spectrum from different pixels. **d**, Overlay image of lipids and protein channel indicates some small dots in a large exosome are rich in lipids. All images were acquired with a 1.05 NA objective in 1× PBS (expansion factor, 4.5). Scale bars, 10 µm in (**a**, **b**); 1 µm in (**d**).

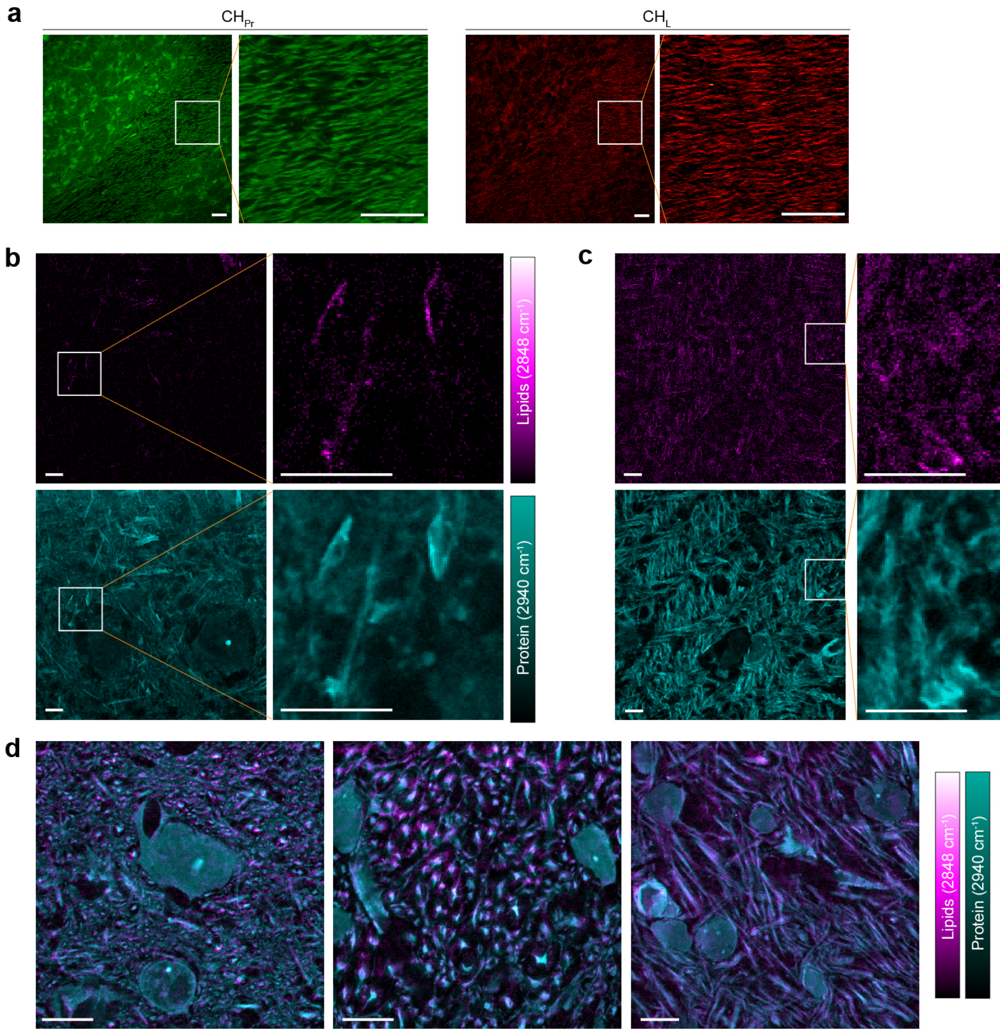


**Supplementary Figure 7. Imaging of protein and lipids in the expanded mouse brain tissue.** Representative protein and lipid channels in expanded mouse brain (a) 2-fold expansion, (b-c) 2.56-fold expansion, (d) 4.88-fold expansion. All images were acquired with a 25× objective. Scale bars, 5 µm in (**b-d**); 10 µm in (**a**).

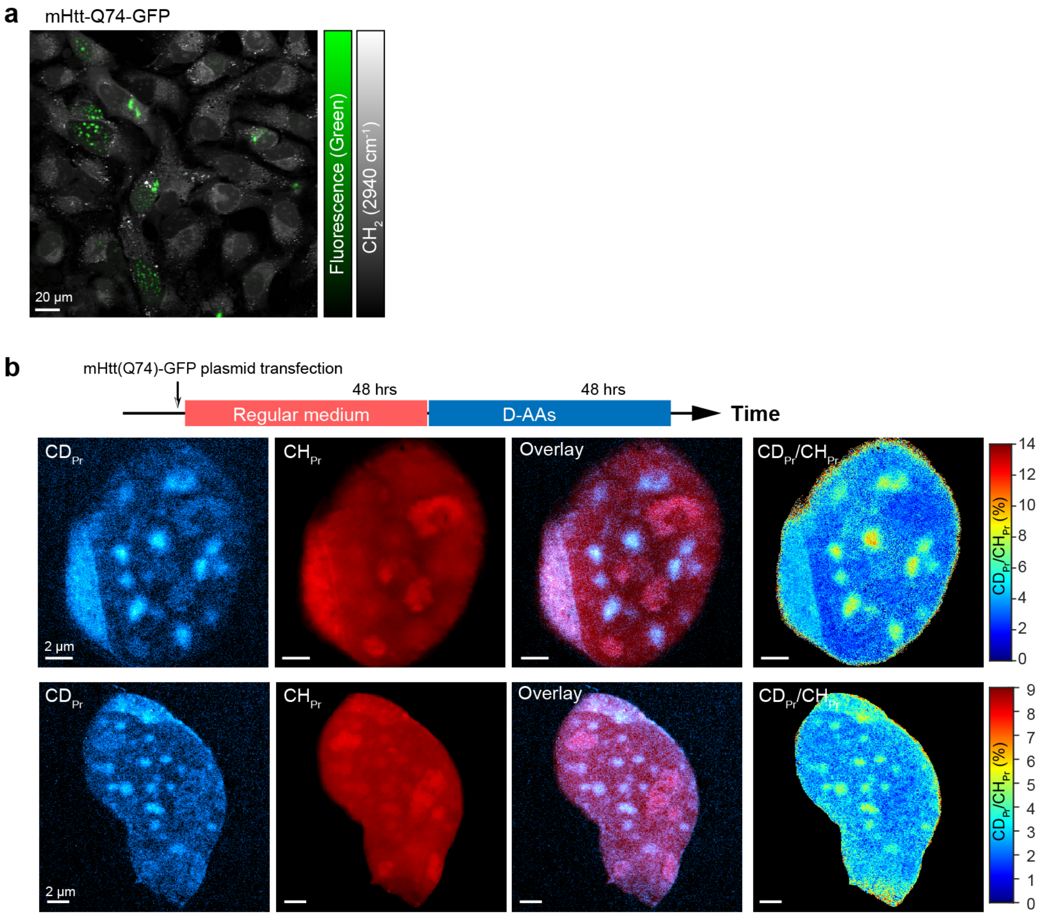


**Supplementary Figure 8. Nanoscale imaging of protein metabolism in Huntingtin aggregates with isotope labeling. a**, Expression of mutant huntingtin polyQ protein was confirmed with both fluorescence (green channel) and SRS ($CH_3$ channel, 2941 $cm^{-1}$) imaging before expansion. Deuterated amino acid labeling in time with simultaneous expression of mutant huntingtin (mHtt74Q-GFP) proteins for 48 hrs. **b**, Regular DEME medium was applied with simultaneous expression of mutant huntingtin (mHtt74Q-GFP) proteins for 48 hrs, chased with deuterated amino acid labeling for another 48 hrs. Images were acquired with a 25× objective in 5× PBS (expansion factor, 3.44). Scale bars, 20 µm in (**a**); 2 µm in (**b**).

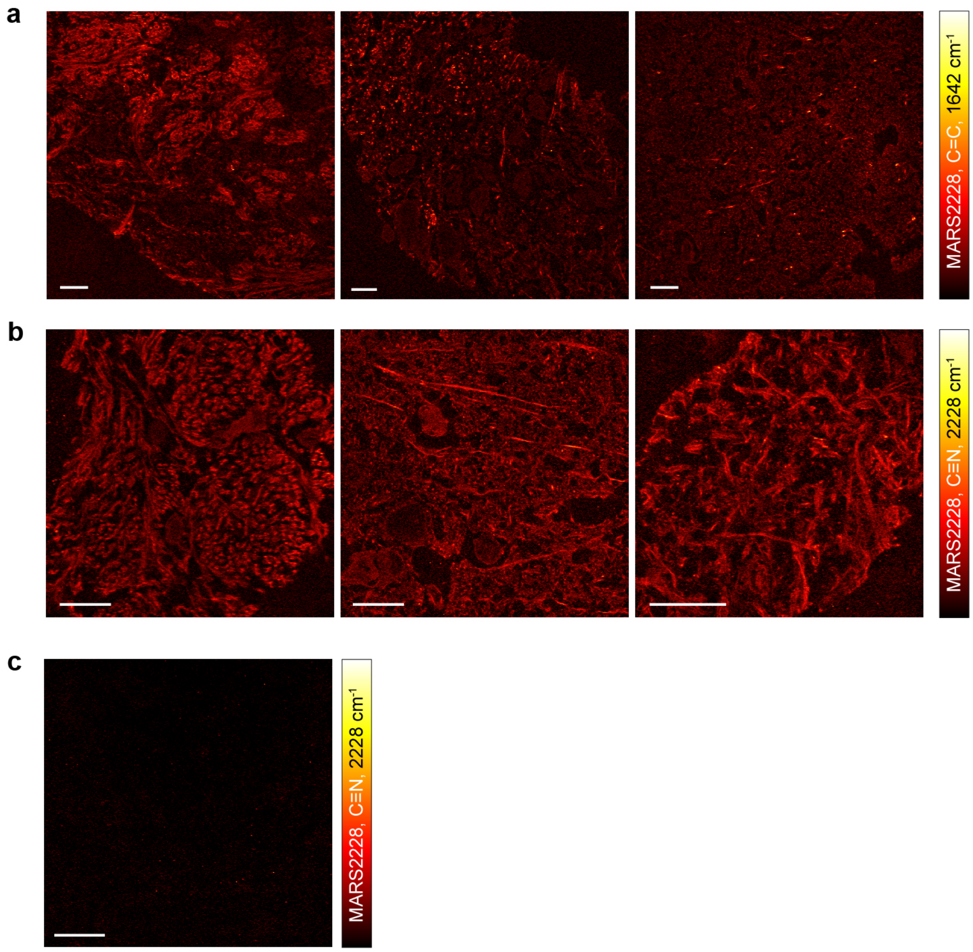


**Supplementary Figure 9. Suppression of non-specific staining of MARS dyes. a-b**, Nonspecific staining backgrounds were observed when 1× PBS was used as the staining buffer. In (**a**), expanded mouse brain tissue was first labeled with rabbit anti-Synaptophysin followed by labeling with donkey-anti-rabbit MARS2228 antibody. SRS images were acquired by probing the C=C bond at 1642 $cm^{-1}$. In (**b**), expanded mouse brain tissue was first stained with mouse anti-$\alpha$ tubulin followed by labeling with donkey-anti-mouse MARS2228 antibody. SRS images were acquired by probing a nitrile bond at 2228 $cm^{-1}$. **c**, Correct staining patterns of Synaptophysin using 9× PBS with 10% Triton-X as the staining buffer. Expanded mouse brain tissues stained with rabbit anti-Synaptophysin primary antibody and donkey-anti-rabbit MARS2228 antibody and imaged via the nitrile bond at 2228 $cm^{-1}$. Scale bars, 10 μm.

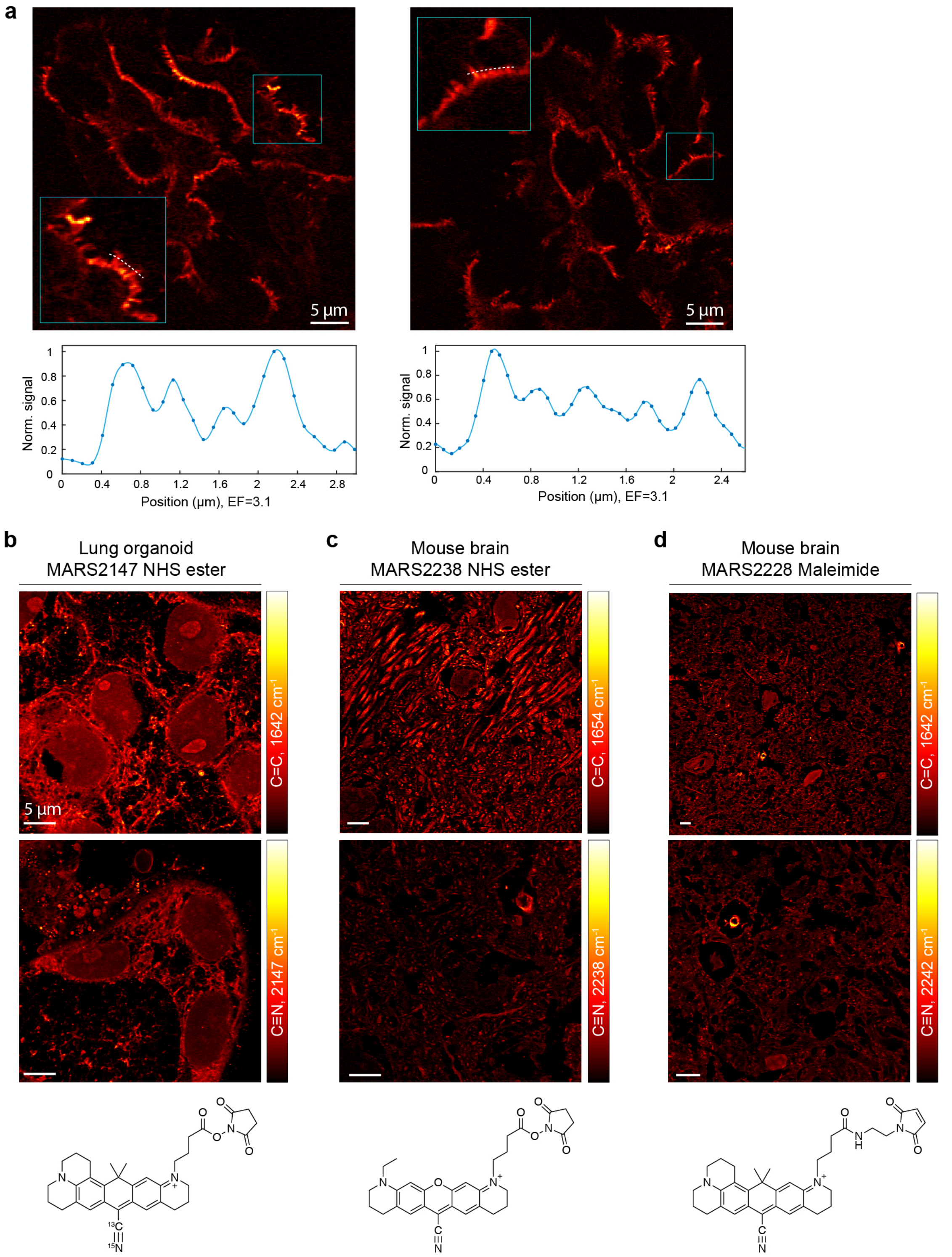


**Supplementary Figure 10. Epr-SRS imaging of immuno-staining and pan-staining with MARS probes. a**, Immuno-eprSRS of actinin-4 (ACTN4, specifically label tertiary podocyte foot processes) in human kidney FFPE tissue with MARS2228. Inset, zoom into the region outlined by the blue box, dotted white curve within the inset indicates the line cut analyzed below. Below, normalized epr-SRS signal along the line cut of the inset. **b-d**, Pan-staining with (**b**) NHS-ester-functionalized MARS2147 on expanded human lung organoid, (**c**) NHS-ester-functionalized MARS2238 on expanded mouse brain tissue and (**d**) Maleimide-functionalized MARS2228 on expanded mouse brain tissue. Chemical structures of MARS probes were shown below images. Images were acquired with a 25× objective. (**a**) were imaged in 1/50 PBS (3.1-fold expansion); (**b-d**) were imaged in 1× PBS (4.5-fold expansion). Scale bars, 5 μm in (**b-d**).

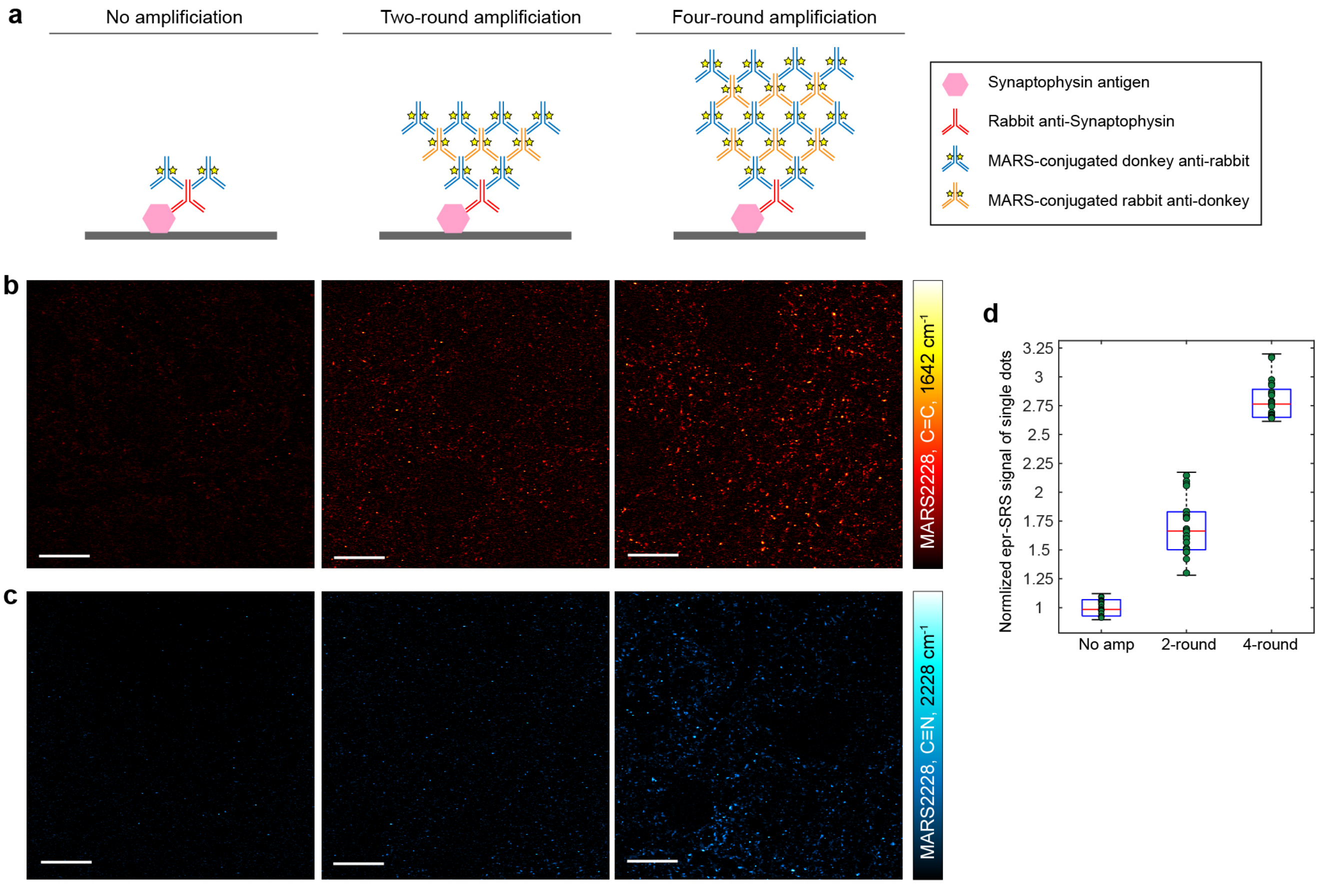


**Supplementary Figure 11. Signal amplification via cyclic staining. a**, Schematic explanation of signal amplification of epr-SRS via cyclic staining of MARS-conjugated secondary antibodies. **b-c**, Epr-SRS images of Synaptophysin in mouse brain via (**b**) C=C bond at 1642 $cm^{-1}$ and (**c**) C≡N bond at 2228 $cm^{-1}$ of MARS2228. Left to right: no amplification, amplification with additional two rounds staining and amplification with additional four rounds staining. **d**, Quantification of amplification factors (mean$\pm$s.d.) with C=C band at 1628 $cm^{-1}$ of MARS2228. Strongest 20 dots were selected for analysis in each condition. Images were acquired with a 25× objective in 1× PBS (4.5-fold expansion). Scale bars, 10 μm.

**Supplementary Table 1. Reagents**

| Name | Source | Catalog No. |
|---|---|---|
| A8301 (inhibitor of transforming growth factor β kinase type 1 receptor) | Sigma-Aldrich | SML0788-5MG |
| BEBM™ Bronchial Epithelial Cell Growth Basal Medium | Lonza | CC-3171 |
| BEGM™ Bronchial Epithelial Cell Growth Medium SingleQuots™ Supplements and Growth Factors | Lonza | CC-4175 |
| Dulbecco's Modified Eagle Medium | Gibco™ | 11965 |
| CHIR99021 (activator of WNT pathway) | Reprocell | 04000402 |
| Dulbecco's Phosphate-Buffered Saline (DPBS), 1X without calcium and magnesium | Corning | 21-031-CV |
| DMH-1 (Inhibitor of BMP4/SMAD signaling) | Tocris Bioscience | 4126 |
| Growth factor reduced Matrigel | Corning | CB 40230 |
| Heparin solution | Stemcell Technologies | 07980 |
| HyClone™ FetalClone™ I Serum (a fetal bovine serum alternative) | GE Healthcare | SH30080.03 |
| Hydrocortisone stock solution | Stemcell Technologies | 07925 |
| Normal Human Bronchial Epithelial (NHBE) cells without Retinoic Acid | Lonza | CC-2541 |
| Paclitaxel | Cayman Chemicals | 10461 |
| PneumaCult™-ALI Basal Medium | Stemcell Technologies | 05002 |
| PneumaCult™-ALI 10X Supplement | Stemcell Technologies | 05003 |
| PneumaCult™-ALI Maintenance Supplement | Stemcell Technologies | 05006 |
| Penicillin-Streptomycin | Lonza Walkersville Inc. | 17-602E |
| RPMI 1640 with L-glutamine | Corning | 10-040-CV |
| Y27632 (Inhibitor of ROCKs) | Cayman Chemical | 129830-38-2 |

| | | |
|---|---|---|
| 0.25% Trypsin-EDTA | Gibco™ | 25200056 |
| Proteinase K | Thermo Scientific | EO0491 |
| Propyl gallate | Sigma Aldrich | P3130 |

**Supplementary Table 2. Calibrated expansion factors of various samples in different buffer conditions**

| Sample type | Measured buffer condition | Expansion factor | | | Calculation method utilized |
|---|---|---|---|---|---|
| | | Mean | STE | N | |
| Mouse kidney | 1× PBS | 4.30 | 0.07 | 15 | Distances |
| | 1:25× PBS | 7.43 | 0.27 | 10 | |
| | 1:50× PBS | 8.10 | 0.19 | 12 | |
| FFPE Human Kidney | 1× PBS | 2.30 | 0.54 | 20 | Average nuclear size |
| | 1:50× PBS | 3.11 | 0.94 | 15 | |
| Mouse brain (ExPath Gel[1]) | 1× PBS | 1.99 | 0.24 | 1724 | |
| | 1:25× PBS | 2.56 | 0.37 | 1062 | |
| HeLa cells | 5× PBS | 3.44 | 0.68 | 110 | |
| Mouse brain (MAGNIFY Gel[2]) | 10× PBS | 2.96 | 0.68 | 82 | |
| | 5× PBS | 3.23 | 0.63 | 189 | |
| | 2× PBS | 3.85 | 0.56 | 414 | |
| | 1× PBS | 4.48 | 0.62 | 403 | |
| | 1:10× PBS | 5.38 | 0.82 | 163 | |
| | 1:50× PBS | 7.18 | 1.22 | 165 | |
| | Water | 10.13 | 1.60 | 133 | |

[1]ExPath Gel formula: 15 g/100mL Sodium Acrylate, 5 g/100 mL Acrylamide, 500 ppm N,N′-Methylenebisacrylamide (Bis), 11.7 g/100 mL NaCl, 1x PBS.
[2]MAGNIFY Gel Formula: 4 g/100 mL N,N-Dimethylacrylamide (DMAA), 34 g/100 mL Sodium Acrylate, 10 g/100 mL Acrylamide, 100 ppm N,N′-Methylenebisacrylamide (Bis), 1 g/100 mL NaCl, 1x PBS.

**Supplementary Table 3. Antibody summary**

| Primary antibodies | | | | | |
|---|---|---|---|---|---|
| Target | Vendor | Catalog # | Host species[a] | Concentration (mg/mL) | Clonality[b] |
| ACTN4 | Sigma | HPA001873 | Rabbit | 0.2 | pAb |
| MAP2 | SYSY | 188004 | Guinea Pig | / | pAb |
| Synaptophysin | Proteintech | 17785-1-AP | Rabbit | / | pAb |
| PSD95 | Abcam | ab12093 | Goat | 1 | pAb |
| TH | Abcam | ab76442 | Chicken | 0.2 | pAb |
| αTubulin | Sigma | T6199 | Mouse | 1 | mAb |
| Vimentin | Abcam | ab24525 | Chicken | / | pAb |

| Secondary antibodies | | |
|---|---|---|
| Target | Source | Catalog # |
| Bovine anti-Goat IgG (H+L) | Jackson ImmunoResearch | 805-005-180 |
| Rat anti-Mouse IgG (H+L) | Jackson ImmunoResearch | 415-005-166 |
| Goat anti-Bovine IgG (H+L) | Jackson ImmunoResearch | 101-005-165 |
| Mouse anti-Rat IgG (H+L) | Jackson ImmunoResearch | 212-005-168 |
| Donkey anti-Rabbit IgG (H+L) | Jackson ImmunoResearch | 711-005-152 |
| Rabbit anti-Donkey IgG (H+L) | Thermo Fisher | SA1-26816 |
| Donkey anti-Guinea Pig IgG (H+L) Alexa Fluro 488 | Jackson ImmunoResearch | 706-545-148 |
| Donkey anti-Chicken IgY (H+L) Cy3 | Jackson ImmunoResearch | 703-165-155 |
| Goat anti-Rabbit IgG (H+L) | Thermo Fisher | SA5-10041 |

| Lectins | | | | |
|---|---|---|---|---|
| Target | Conjugate | Source | Catalog # | Concentration (mg/mL) |
| *Lycopersicon Esculentum* lectin | DyLight488 | Vector | DL-1174 | 1 |
| *Lycopersicon Esculentum* lectin | unconjugated | Vector | L-1170 | / |
| Wheat Germ Agglutinin | unconjugated | Sigma Aldrich | L0636 | / |

| Non-protein based | | |
|---|---|---|
| Name | Source | Catalog # |
| NucBlue Fixed Cell | Invitrogen | R37606 |
| DAPI | Thermo Fisher | 62248 |

**Supplementary Video 1.** 3D-rendered SRS images of $CH_3$ peak at 2941 $cm^{-1}$ of an expanded mouse brain tissue (2-fold expansion) taken at 25× magnification.

**Supplementary Video 2.** 3D-rendered SRS images of $CH_3$ peak at 2941 $cm^{-1}$ of an expanded human lung organoid (4.5-fold expansion) taken at 25× magnification.

**Supplementary Video 3.** 3D-rendered epr-SRS image of MARS2228-conjugated *Lycopersicon Esculentum* lectin (LEL) labeling (red) and $CH_3$ image of 2941 $cm^{-1}$ (green) of an expanded mouse brain tissue (4.5-fold expansion) taken at 25× magnification.

**Supplementary Video 4.** 3D-rendered epr-SRS image of expanded human lung organoid (4.5-fold expansion) stained with MARS2147 NHS ester dye taken at 25× magnification.

**Supplementary Video 5.** 3D-rendered epr-SRS image of MARS probe immunolabeled Synaptophysin (green) and PSD95 (red) of an expanded mouse brain tissue (4.5-fold expansion) taken at 25× magnification.
